# Effects of Feed Processing Type, Protein Source, and Environmental Salinity on *Litopenaeus vannamei* Feeding Behaviour

**DOI:** 10.1101/2024.03.14.584959

**Authors:** Amina S. Moss, Adam J. Brooker, Stella N. Ozioko, Marit A. J. Nederlof, Sudip Debnath, Johan Schrama

**Affiliations:** Institute of Aquaculture, University of Stirling, Scotland, United Kingdom; Department of Animal Sciences, Aquaculture and Fisheries Group, Wageningen University and Research, Wageningen, The Netherlands

**Keywords:** *Litopenaeus vannamei*, extruded feed, steamed pellets, growth performance, shrimp feeding behaviour, salinity

## Abstract

In an 8-week trial, the effects of feed processing types (extruded vs. steamed pellets) and protein source (soya/pea vs. fish meal) on *Litopenaeus vannamei* were studied under 30 ppt (first 30 days) and 5 ppt (last 15 days) salinity conditions. Diets included D1-SE soya/pea extruded, D2-SS soya/pea steamed, D3-FE fishmeal extruded, and D4 (FS) fishmeal steamed pellets. No significant weight differences were observed. Survival rates (80-97% at 30 ppt) decreased at reduced salinity and were significantly higher for shrimp fed steamed pellets (D2:80.00%, D4:76.67%) compared to extruded (D1: 50.00%, p<0.05). Shrimp fed D2-SS and D4-FS diets had increased lipid content (6.79%, 6.36%, p<0.05). Significantly lower lysine and isoleucine were noted in D2-SS. Behaviourally, at 30 ppt, D3-FE attracted significantly more shrimp (84.72%) than D1-SE (74.49%) and D2-SS (75.70%, p<0.05). Click analysis showed D1-SE and D2-SS with the shortest durations (17.97, 17.24ms, respectively), and D3-FE and D4-FS the longest (20.10, 23.89ms, respectively, p<0.05). Click frequency was also significantly higher in fishmeal–based diets, whereas the number of clicks was higher in shrimp fed extruded feed. Overall, extruded pellets and fishmeal diets were more favourable. These findings emphasize the importance of tailored feed strategies that consider nutritional content, feed physical properties and environmental factors for optimal shrimp feeding.

## 1. Introduction

Understanding the feeding behaviours of key aquaculture species, such as the Pacific white shrimp (*Litopenaeus vannamei*), is essential not only for advancing global shrimp production but also for addressing ecological concerns. Dominating the crustacean farming sector, this species accounts for approximately 80% of worldwide shrimp cultivation (FAO 2022). The last two decades have witnessed a remarkable expansion in shrimp aquaculture. From 2000 to 2018, production increased fivefold, reaching a staggering six million tons (FIGIS 2018). Despite this success, the industry faces significant challenges, particularly in feeding practices, which are critical both economically and environmentally. Inefficient feeding strategies contribute to over half of the total production costs and lead to substantial losses (Silva et al. 2012; Tacon et al. 2013). Moreover, the slow feeding response of shrimp and consequent nutrient discharge of uneaten feeds into water bodies present serious environmental concerns (Liao et al., 2011; Iber et al, 2021; Quiroz-Guzman et al., 2022).

In addressing these issues, studying shrimp feeding behaviour is essential. As benthic feeders, shrimp rely heavily on chemoreceptors for food detection due to their underdeveloped eyesight (Obaldo et al. 2006). Their feeding is mediated by chemoreceptors located on antennules, antennae, mouthparts, and legs, which are essential in efficiently locating and consuming food to prevent nutrient leaching from the feed (Gadient & Schai 1994). This unique feeding behaviour, combined with their limited digestive capacity, requires continuous consumption of small feed quantities, often resulting in significant feed wastage and water quality degradation in holding ponds (Sanchez et al. 2005; Jescovitch et al. 2018; Reis et al. 2020). The challenge, therefore, is to develop feeding strategies that maximize efficiency while minimizing waste and environmental impact.

Given these complexities, there has been a growing emphasis on in-depth research into shrimp feeding behaviours to enhance aquaculture practices and shrimp welfare. Studies have demonstrated that monitoring behavioural patterns is vital for identifying stress or disease indicators in shrimp, necessitating intensive observation due to their intricate feeding habits (Costa et al. 2016; Bardera et al. 2019, 2020). The physical process of shrimp feeding involves the use of chelate pereiopods for food capture and transportation towards the mouthparts. The coordination of various mouthparts, such as mandibles, maxillipeds, and maxillae, is critical in the feeding process (Smith and Shahriar 2013). Furthermore, shrimp have been found to produce clicking sounds. These sounds, believed to be a result of rapid closure of shrimp’s mouthparts during feeding (Nguyen et al., 2018), may be influenced by the texture and hardness of feed particles. Factors influencing shrimp feeding behaviour include physiological changes, feed deprivation, moulting, stocking densities, and environmental conditions like salinity and feed types (Bardera et al. 2019).

The study of feed composition reveals that ingredients and chemical compounds in feed play a significant role in triggering various stages of feeding behaviour. Commercial feeds are designed with specific chemical cues for rapid food recognition by shrimp (Lee and Meyers 1996; Sanchez et al. 2005). The use of diverse protein sources, both terrestrial and marine, in shrimp feed production has been extensively researched (Samocha et al. 2004; Nunes et al. 2006; Tantikitti 2014; Galkanda-Arachchige et al. 2019; Galkanda- Arachchige and Davis 2020). The exploration of alternative protein sources, particularly plant–based proteins like soybean meal, has been a focal point due to the high costs associated with fishmeal (Samocha et al. 2004). This transition is not without challenges, as alternative protein sources often require supplementation due to differences in amino acid profiles (Galkanda et al., 2019; Galkanda et al., 2020; Howlader et al., 2023; Nunes et al., 2023). Moreover, the processing methods of feed pellets, such as extrusion and steam pelleting, significantly affect their nutritional value, digestibility, and environmental impact, necessitating a deeper understanding of these methods for effective shrimp feed management (Hilton et al., 1981; Booth et al., 2002; Zhou, 2014; Hoyos et al., 2017; Gao et al., 2019; Aaqillah-Amr, 2021; Espinoza-Ortega et al., 2023).

Furthermore, environmental factors, particularly salinity, play a crucial role in influencing shrimp physiology and behavior. As climate change affects salinity patterns globally, understanding its impact on shrimp is essential for developing adaptive feeding strategies. Studies have shown that water salinity and dietary protein content significantly affect the growth performance of *L. vannamei* juveniles, with better performance observed at higher salinity and protein levels (Bardera et al., 2019; Rahi et al. 2021; Pinho and Emerenciano, 2021; Ballantyne et al., 2023;). While the long-term impacts of salinity on shrimp have been explored, the immediate effects on feeding behaviour and how it interacts with environmental conditions remain under-studied.

To comprehensively address these multifaceted challenges, this study uses Quantitative Shrimp Feeding Behavioural Assessment (QSFBA) based on methodologies in past behaviour studies (Wemelsfelder, 2007; Barnard et al., 2016; Darodes de Tailly et al., 2021) by employing techniques such as real-time image segmentation and Passive Acoustic Monitoring (PAM) for observing and analysing shrimp feeding behaviour. These non-invasive methods provide valuable insights into the intricate patterns of shrimp feeding in response to different feed types and environmental factors (Jescovitch et al. 2018; Darodes et al. 2021; Smith & Shahriar 2013; Smith & Tabrett 2013; Peixoto et al. 2020). The current study, therefore, investigates the nutritional profile, growth performance, and feeding behaviour of *L. vannamei* juveniles in relation to feed types (steamed vs. extruded pellets), protein sources (soya/pea vs. fishmeal), and varying salinity levels (5 ppt and 30 ppt). By using video recordings and PAM, the study aims to understand their feeding patterns and growth under these conditions. The ultimate goal is to use an interdisciplinary approach to optimize feeding strategies in shrimp farming, thus enhancing both the economic efficiency and environmental sustainability of this vital industry and contributing to the field of aquaculture research.

## 2. Materials and methods

The animal experiments in this study were conducted in accordance with the ethical guidelines and approved by the ethics committee of the University of Stirling (AWERB 2022 10523 7859).

### 2.1. Diet preparation

Four isonitrogenous (36.98 ± 0.66%) and isolipidic (7.58 ± 0.34%) experimental diets were prepared at Wageningen University (Netherlands) to assess shrimp growth performance and feeding behaviour as follows: D1 SE (soya and pea protein, extruded feed), D2 SS (soya and pea protein, steamed pellets), D3 FE (fishmeal protein, extruded feed) and D4 FS (fishmeal protein, steamed pellets), where soya and pea proteins are added 1:1. The composition and proximate analysis of these diets are detailed in Table 1. These diets were formulated to meet the nutritional requirements for shrimp as per previous literature (Wouters et al. 2001; Moss et al., 2019). The main protein sources used were pea protein, soya protein concentrate and fishmeal. Dry ingredients were first weighed and thoroughly mixed using a mechanical mixer to ensure a homogeneous blend. Then liquid ingredients were added, and the diet mixed again. For diets that required extrusion, the mixture was processed using an extruder (Clextral BC45 laboratory scale twin-screw) under controlled temperature and pressure conditions. The extruded diets were then cooled, dried, and stored in a freezer (−20°C). For diets that required steaming, the mixture was steamed ensuring proper cooking to enhance digestibility. The steamed diets were subsequently cooled, dried, and stored in a freezer (−20°C). The prepared diets were analysed by the Nutrition Analytical Services laboratories (University of Stirling, Scotland) for proximate composition.

**Table 1.** Formulation and proximate analysis of the experimental diets (% dry matter).

| Diets | D1 SE (Soya/Pea Extruded) | D2 SS (Soya/Pea Steamed) | D3 FE (Fishmeal Extruded) | D4 FS (Fishmeal Steamed) |
| --- | --- | --- | --- | --- |
| Processing Method | Extruded | Steamed | Extruded | Steamed |
| **Ingredients** |  |  |  |  |
| Wheat | 18.05 | 18.05 | 20.60 | 20.60 |
| Wheat Flour (Native) | 17.00 | 17.00 | 20.00 | 20.00 |
| Wheat Gluten | 10.00 | 10.00 | 10.00 | 10.00 |
| Wheat Bran | 15.00 | 15.00 | 15.00 | 15.00 |
| Soya Protein Concentrate | 15.00 | 15.00 | --- | --- |
| Pea Protein Concentrate | 15.00 | 15.00 | --- | --- |
| Fishmeal LT (CP>680) | --- | --- | 30.00 | 30.00 |
| Soya Lecithin | 0.70 | 0.70 | 0.70 | 0.70 |
| Fish Oil | 3.00 | 3.00 | 1.00 | 1.00 |
| Cholesterol | 0.30 | 0.30 | 0.30 | 0.30 |
| CaCO<sub>3</sub> | 1.60 | 1.60 | 0.30 | 0.30 |
| Mono Calcium Phosphate | 2.20 | 2.20 | 0.50 | 0.50 |
| L-Lysine | 0.35 | 0.35 | 0.20 | 0.20 |
| DL-Methionine | 0.55 | 0.55 | 0.20 | 0.20 |
| L-Threonine | 0.25 | 0.25 | 0.20 | 0.20 |
| Premix | 1.00 | 1.00 | 1.00 | 1.00 |
| **Total** | **100.00** | **100.00** | **100.00** | **100.00** |
| **Proximate Analysis** |  |  |  |  |
| Moisture (%) | 5.28 | 5.89 | 4.18 | 6.60 |
| Lipid (%) | 8.00 | 7.34 | 7.71 | 7.28 |
| Crude Protein (%) | 37.31 | 36.34 | 37.74 | 36.52 |
| Ash (%) | 5.88 | 5.93 | 7.08 | 6.93 |
| Energy (kJ/g) | 19.25 | 18.92 | 19.07 | 18.66 |
| TBARS (mg/kg) | 3.21 | 3.14 | 5.76 | 5.33 |

### 2.2. Feeding trial

The feeding trial was conducted over 8 weeks in 100L clear glass aquarium tanks in a recirculation system at Wageningen University. Constant aeration was supplied, and the water quality parameters were maintained at pH 8.2 ± 0.4 (mean ± SD), temperature 25– 28.0°C, dissolved oxygen 7.3 ± 0.8 mg/L, ammonia 0.1 ± 0.15 mg/L, and nitrate 0–75 mg/L. In this feeding trial, a deliberate alteration in salinity was employed as a key variable to assess its impact on shrimp at different life stages. The trial was initially conducted at a salinity of 30 ppt for the first 45 days, catering to the younger juvenile shrimp. Subsequently, for the final 15 days, the salinity was reduced to 5ppt, a shift designed to examine the adaptability and feeding behaviour of the shrimp as they progressed to a slightly more mature stage. This approach allowed for a better insight into the effects of salinity changes on the growth of early juvenile shrimp as they mature.

Ten shrimp (initial wet weight 0.53**–**0.56 g per individual) were randomly stocked in each tank. Each of the four treatments was tested in triplicate (n = 12 tanks). Individual tank dimensions were 90 cm × 45 cm X 20 cm. The daily husbandry protocol involved siphoning faeces, moulted exoskeleton and uneaten feeds followed by the setup of Go- Pro cameras (GoPro Hero9, GoPro Inc., San Mateo, CA, USA), hydrophones (Aquarian Hydrophone H2D, Aquarian Audio & Scientific, Anacortes, WA, USA) and recorders (Zoom H5 Handy Recorder, Sound Service MSL, Reading, UK), then feeding. Daily feed provided was 5% of each tank’s shrimp biomass. Shrimp were counted and weighed weekly to adjust feed inputs.

Growth performance and survival outcomes of *L. vannamei* shrimp were evaluated as follows:

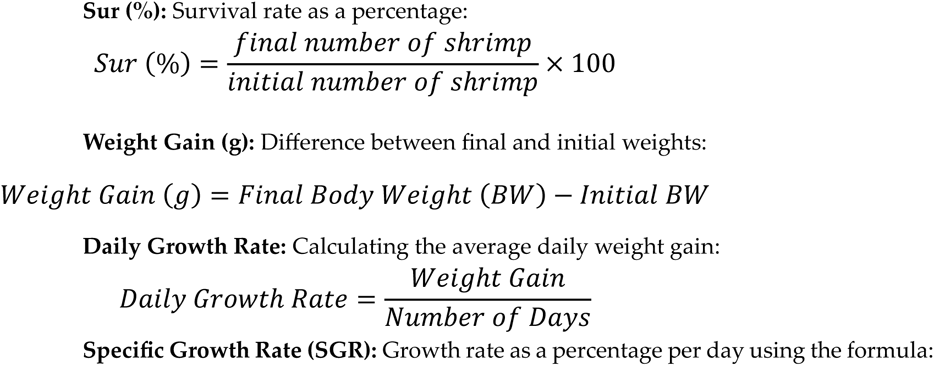

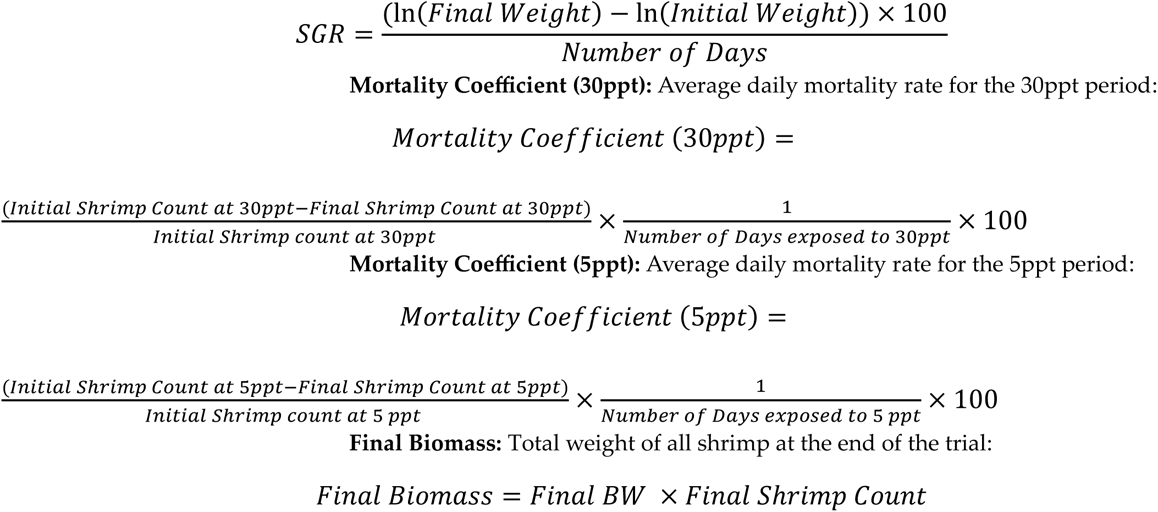

At the end of the feeding trial, shrimp were euthanized and collected for biochemical analysis. These samples were stored in a freezer (−20°C) until they were freeze-dried.

### 2.3. Biochemical composition

The analyses of moisture (AOAC, 1995), crude protein (AOAC, 1995), ash (AOAC, 1995) and crude lipid (Folch et al., 1957) composition of whole shrimp and experimental diets were conducted according to the respective citations. To obtain fatty acid methyl esters, the crude lipids were transmethylated using the procedure outlined by Christie (1982). The resulting esters were then separated via gas−liquid chromatography (GLC) and quantified using a flame ionization detector (FID) on a GC-14A instrument (Shimadzu, Tokyo, Japan) following the conditions described by Izquierdo et al. (1990). Identification of the fatty acids was achieved through gas-liquid chromatography-mass spectrometry (GLC-MS) and comparing the samples with previously characterized standards. The levels of lipid oxidation in the feeds were assessed by the 2-Thiobarbituric Acid Reactive Substances (TBARS) assay according to methods described by Pfalzgraf (1995). Total amino acid concentrations in feeds and whole shrimp were determined using High-Performance Liquid Chromatography (HPLC; Shimadzu Corp., Tokyo, Japan) as described by Kader et al. (2010).

### 2.4. Feeding behaviour analysis

#### 2.4.1. Data Collection and Equipment Setup

To assess shrimp feeding behaviour, the current study used Quantitative Shrimp Feeding Behavioural Assessment (QSFBA) by employing imaging data and acoustics, based on methodologies in animal behaviour studies (Wemelsfelder, 2007; Barnard et al., 2016; Darodes de Tailly et al., 2021; Barreto et al., 2022). During the 8-week feeding trial, shrimp were video recorded interacting with the provided feed once per day for the last 33 days. Four GoPro cameras were used to record four tanks each day, one from each treatment, with the recorded replicate changing each day so that each tank was recorded every third day. The GoPro cameras were positioned above the tank facing downwards in a hole cut in the centre of the tank lid. The cameras were set to capture high-resolution images every second for a 60-minute period prior to and following the introduction of the feed. These observations were conducted consistently at the same hour each day to maintain a uniform 23-hour feed deprivation period throughout the study, as previously outlined by Bardera et al. (2020). Additionally, two hydrophones and recorders were used to record sound continuously for one hour immediately after feed was dispensed in the tanks concurrently with two of the cameras with each tank recorded every sixth day.

#### 2.4.2. Image Analysis using Image J Software Application

The recorded images were processed and analysed using Image J Software (Fiji version 2.9.0) according to procedures previously explained (Rolston, 1995). The software’s sequence package facilitated the stacking of images into video format, which enhanced contrast and threshold settings, thereby rendering the shrimps and feed pellets prominently visible. To track the movement of each shrimp within each replicate group in real-time during recordings, the software’s built-in manual tracking tool and timer were employed. The behaviours measured were categorized according to previous studies (Pontes, 2008; Bardera et al., 2020; Peixoto et al., 2020) to assess the following feeding behaviour metrics: attraction to feed (based on latency), feeding time, zone transitions, average distance to feed, feed intake, shrimp length, full/empty gut, and growth rate (Table 2; Figure 6).

**Figure 1.**
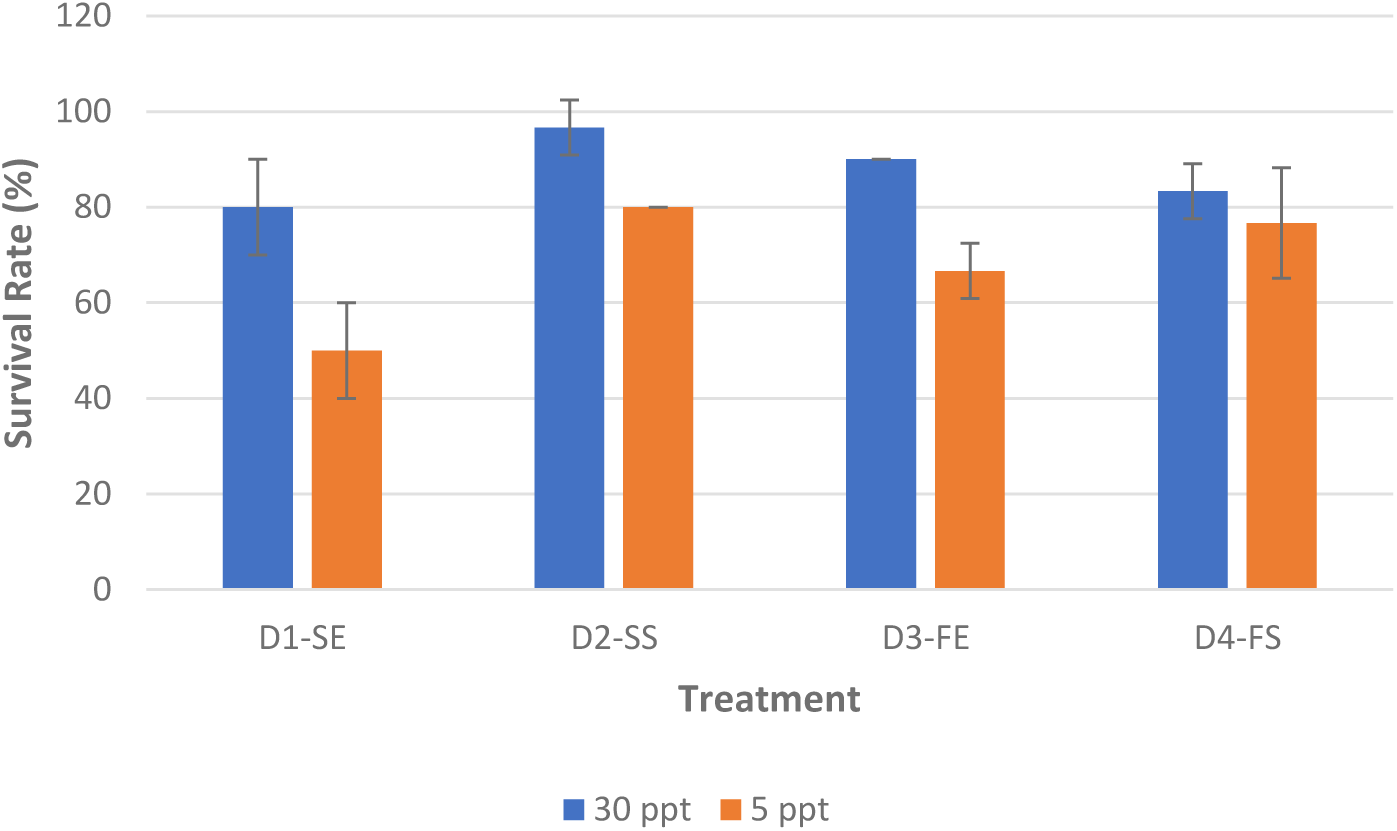
Survival Rates Across Two Salinity Conditions.

**Table 2.** Feeding Behaviour parameters of *L. vannamei* measured using Image J Software Application (Fiji version 2.9.0).

| Behaviour | Description | Formula |
| --- | --- | --- |
| Attraction –to–feed (%) | Time (% of total recording time) to reach the feeding area Inverted results were used to represent attraction rather than latency. | $100 - \left( \frac{3600 - \text{Time taken to reach feeding area}}{3600} \times 100 \right)$ |
| Feeding time (min) | Time (min) on the feeding area consuming food. | No formula |
| Transitions | Count of shrimp entering the feeding area. | Number of transitions |
| Distance–to–feed (cm) | Average distance (cm) between shrimp and the feeding area. | No formula |
| Feed intake (%) | Percentage of feeding pellets consumed | $\frac{(\text{Initial pellets} - \text{Final Pellets})}{\text{Initial pellets}} \times 100$ |
| Mean shrimp length (cm) | Mean shrimp length on the initial and final days (cm) | No formula |
| Full gut / empty gut ratio | The ratio of shrimp with full gut vs. shrimp with empty gut one hour after feed dispensed | $\frac{\text{Number of shrimp with full gut}}{\text{Number of shrimp with empty gut}}$ |
| Daily growth rate (cm/day) | Daily growth rate calculated from initial and final shrimp lengths | $\frac{\text{Final length} - \text{Initial length}}{\text{Number of days}}$ |

#### 2.4.3. Acoustic Data Analysis

Audio recordings from each experimental diet group were transferred from the hydrophone recorder to a high-performance Windows PC for analysis. The original acoustic data were partitioned into 15-minute intervals to reduce the size of each file. Using MATLAB software version R2023a, a Fast Fourier Transform (FFT) was used to transform the acoustic signals into their constituent frequency components and corresponding amplitudes.

##### Segmentation of Clicking Sounds

Segmenting shrimp clicking sounds involved three signal processing steps to accurately identify and extract clicking sound events:

**Step 1: Framing the Signal**

Continuous signals were divided into smaller, manageable segments using rectangular windows. Each segment comprised 500 consecutive samples extracted from the original signal to allow the signals to be assessed within shorter time intervals, facilitating detailed analysis.

**Step 2: Calculating RMS and Creating an Envelope**

The Root Mean Square (RMS) of each framed segment was calculated, using the formula (Novotny et al., 2008):

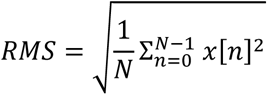

where:

*N* is the number of samples in the window

The sum Σiterates over all data points from *n* = 0 to *N* – 1

The term x[*n*]^2^ represents the square of the data point value at index

The RMS is an indicator of energy or amplitude within a particular segment. The RMS values were used to create an envelope that presents a smoothed rendition of the signal. This envelope accentuates amplitude variations within the signal to aid clicking sound detection and extraction (Yang, 2017).

**Step 3: Clicking Sound Detection and Extraction**

The start of clicking sound events were identified as an RMS values ≥3. A threshold of 3 identified segments with relatively high energy levels, indicative of clicking sounds. The end of clicking sound events were identified where RMS values dropped below 1.5, indicating the conclusion of a clicking sound event. The duration of each clicking sound event was measured in milliseconds (ms) by calculating the time difference between the start and end points in the signal. Energy (in dB) was calculated for each clicking sound event to quantify its acoustic intensity.

Clicking sound components were extracted by generating a detection signal. Initially set at zero, this detection signal becomes active when RMS ≥3 and remains active until RMS <1.5. An element-wise multiplication process isolated the clicking sound segments while attenuating non-clicking segments. The resultant signal predominantly comprised successfully detected clicking sound events and segmented through the utilization of RMS–based thresholding.

##### Signal Preprocessing Using a Band Pass Filter

The original recordings were impeded by background noise, which obscured the clicking sounds emitted by the shrimp. However, frequency analysis revealed that the shrimp clicking sounds spanned a frequency range of 15 kHz to 2 kHz (Peixoto et al., 2020). Consequently, each signal segment was processed using a 4th order Infinite Impulse Response (IIR) band-pass filter to diminish background noise and enhance the clarity of the clicking sounds. The hydrophone signals were subjected to an IIR band-pass filter fashioned using the Butterworth window technique within the frequency range of 15 kHz to 25 kHz (Butterworth, 1930; Smith, 2003). This specific frequency range was chosen subsequent to a comprehensive frequency domain analysis of the initial raw signals.

### 2.5. Statistical analysis

Analysis of variance (ANOVA) with Tukey’s HSD post-hoc test was used to determine significant differences in variables (p <0.05) between the four diets. Data are presented as mean ± standard deviation (SD) or as otherwise specified. For the image data, each behaviour parameter was analysed for potential differences between diets using a one-way ANOVA to assess each diet individually and a two-way ANOVA to assess the influence of feed processing type and protein source on the diets. Two-way ANOVA was also used for nutritional analysis. As for the acoustic data, due to the uneven number of click sounds within each treatment a non-parametric Welch’s ANOVA and Omega squared (ω^2^), as an adjusted R², was used to determine the effect size between treatments. ω^2^ represents the proportion of variance in the data that is explained by the treatment factor.

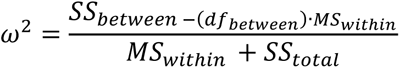

where:

*ω*^2^ represents the adjusted coefficient of determination for one-way ANOVA

*SS*_between_ is the sum of squares between groups

(*df*_between_) is the degrees of freedom associated with between-group variability

*MS*_within_ is the mean square within groups (residual mean square)

*SS*_total_ is the total sum of squares.

According to Field (2013), the thresholds for *ω*^2^ are as follows:

Very small effect size: ES < 0.01

Small effect size: 0.01 <= ES < 0.06

Medium effect size: 0.06 <= ES < 0.14

Large effect size: ES >= 0.14

## 3. Results

### 3.1. Feed composition

Table 1 shows the proximate analysis of experimental diets. Moisture content varied, with Diet D3, FE (Fishmeal protein, extruded feed) at 4.18% (lowest) and Diet D4, FS (Fishmeal protein, steamed pellets) at 6.6% (highest). Lipid content ranged from 7.28% in Diet D4 FS to 8.00% in Diet D1, SE (Soya/pea protein, extruded feed). Minor variations were observed in crude protein content, between 36.34% in Diet D2, SS (Soya/pea protein, steamed pellets) and 37.74% in D3 (FE). TBARS levels were higher in Diets D3 (FE) and D4 (FS) (5.76 mg/kg and 5.33 mg/kg) compared to Diets D1 (SE) and D2 (SS) (3.21 mg/kg and 3.14 mg/kg).

Amino acid composition, detailed in Table S1 (Supplementary Materials), showed that Diet D3 (FE) had higher levels of proline and glycine. The fatty acid profile in Table S2 (Supplementary Materials) indicates that fishmeal–based feeds (Diets D3 and D4) had higher SFAs and MUFAs, while plant–based feeds (Diets D1 and D2) had higher n-6 PUFAs. Diets D3 and D4 were also higher in total n-3 PUFAs. The n-3/n-6 ratio was highest in Diets D3 and D4 (both at 0.57), compared to lower ratios in Diets D1 (SE) and D2 (SS) (0.39 and 0.38, respectively).

### 3.2. Growth performance and proximate analysis of the Pacific white shrimp

The growth performance (Table 3) and nutritional composition (Table 4) of juvenile Pacific white shrimp fed various diets over 8 weeks showed notable differences. Initial body weights were similar across all groups, but final body weight differences, though numerically higher in D1 (SE) and D3 (FE), were not statistically significant across all groups. Survival rates varied significantly across diets, especially at reduced salinity levels, with diets D2 (SS) and D4 (FS) showing higher survival rates compared to D1 (SE) (p<0.05). Feed processing type significantly influenced survival rates and mortality (p<0.05), but protein source did not (p>0.05). Similarly, D1 (SE) was significantly higher than the other diets in terms of mortality coefficient (Figure 2) and daily mortality rate for the 15 days that shrimp were exposed to 5 ppt after being exposed to 30ppt for 45 days prior (p<0.05). Shrimp lipid content differed significantly between diets, with D4 (FS) and D2 (SS) showing higher lipid levels than D3 (p<0.05). Feed processing type affected lipid content (p<0.05), but protein source did not (p>0.05).

**Figure 2.**
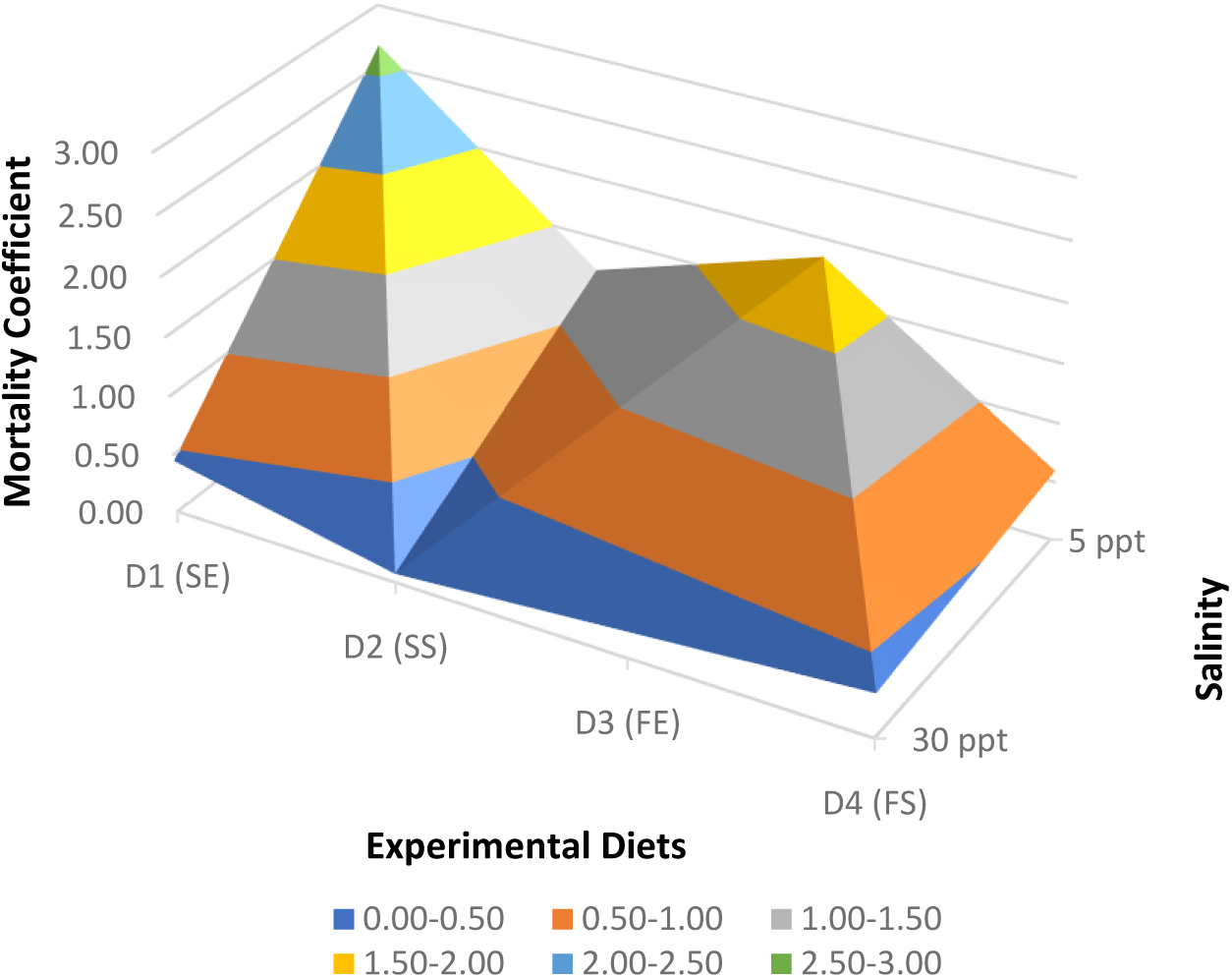
Mortality Coefficient Analysis Across Salinity Conditions. Mortality coefficient of shrimp between the four experimental diets (D1 (SE) = soya and pea extruded, D2 (SS) = soya and pea steamed, D3 (FE) fishmeal extruded and D4 (FS) fishmeal steamed) and two salinity conditions.

**Table 3.** Growth performance of *L. vannamei* fed the experimental diets for 8 weeks. D1 (SE) = soya and pea extruded, D2 (SS) = soya and pea steamed, D3 (FE) = Fishmeal extruded and D4 (FS) = fishmeal steamed.

| Parameter | Test Diets |  |  |  | One-way ANOVA | Two-way ANOVA p-value |  |  |
| --- | --- | --- | --- | --- | --- | --- | --- | --- |
|  | D1 (SE) | D2 (SS) | D3 (FE) | D4 (FS) | p-value | Feed processing type | Protein source | Interaction |
| Initial BW (g)<sup>1</sup> | 0.54±0.06 | 0.54±0.02 | 0.56±0.01 | 0.53±0.04 | 0.800 | 0.503 | 0.880 | 0.503 |
| Final BW (g)<sup>2</sup> | 1.57±0.31 | 1.20±0.11 | 1.39±0.26 | 1.28±0.43 | 0.505 | 0.208 | 0.787 | 0.465 |
| Sur (%)<sup>3</sup> | 50.00±10.00<sup>a</sup> | 80.00±0.00<sup>c</sup> | 66.67±5.77<sup>b</sup> | 76.67±11.55<sup>bc</sup> | 0.008 | 0.003 | 0.195 | 0.067 |
| Weight Gain (g)<sup>4</sup> | 1.03±0.26 | 0.66±0.13 | 0.83±0.26 | 0.75±0.39 | 0.448 | 0.198 | 0.752 | 0.378 |
| Specific Growth Rate (SGR)<sup>5</sup> | 1.48±0.08 | 1.22±0.15 | 1.30±0.20 | 1.29±0.020 | 0.308 | 0.192 | 0.576 | 0.208 |
| Daily Growth Rate (g/d)<sup>6</sup> | 0.02±0.00 | 0.01±0.00 | 0.01±0.00 | 0.01±0.01 | 0.448 | 0.198 | 0.752 | 0.378 |
| Mortality coefficient (30ppt)<sup>7</sup> | 0.44±0.22 | 0.07±0.13 | 0.22±0.00 | 0.37±0.13 | 0.054 | 0.217 | 0.667 | 0.014 |
| Mortality coefficient (5ppt)<sup>8</sup> | 2.66±0.80<sup>c</sup> | 1.22±0.37<sup>ab</sup> | 1.85±0.46<sup>bc</sup> | 0.60±0.52<sup>a</sup> | 0.010 | 0.003 | 0.057 | 0.776 |
| Daily Mortality Rate (30ppt)<sup>9</sup> | 4.44±2.22 | 0.74±1.28 | 2.22±0.00 | 3.70±1.28 | 0.054 | 0.217 | 0.667 | 0.014 |
| Daily Mortality Rate (5ppt)<sup>10</sup> | 21.43±7.14<sup>c</sup> | 11.90±4.12<sup>ab</sup> | 16.67±4.12<sup>bc</sup> | 4.76±4.12<sup>a</sup> | 0.020 | 0.006 | 0.076 | 0.694 |
| Final Biomass (g)<sup>11</sup> | 8.05±3.19 | 9.60±0.90 | 9.22±1.63 | 9.78±3.03 | 0.812 | 0.465 | 0.638 | 0.731 |

Values are means of triplicate groups and presented as mean ± SD. Values in the same row having different superscript letters are significantly different (p < 0.05). The lack of superscript letter indicates no significant differences among treatments.

**Table 4.**
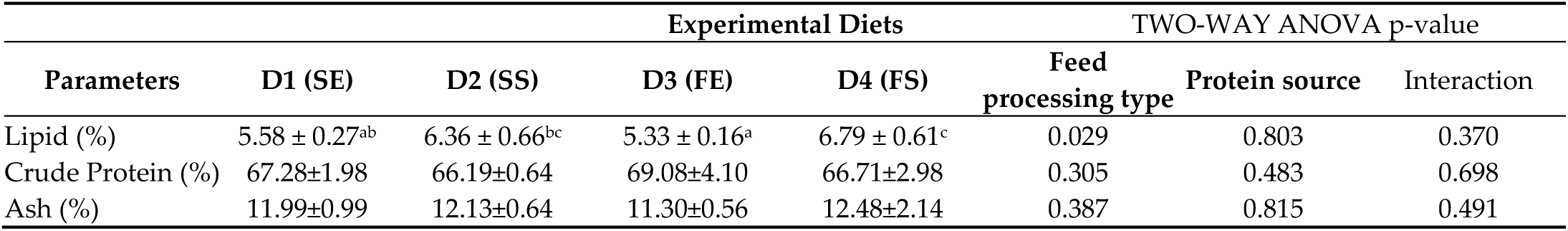
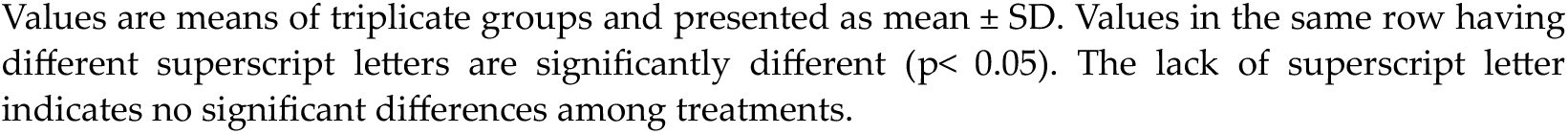
Proximate analysis (% wet weight) of *L. vannamei* fed after being fed the experimental diets for 8 weeks. D1 (SE) = soya and pea extruded, D2 (SS) = soya and pea steamed, D3 (FE) = Fishmeal extruded and D4 (FS) = fishmeal steamed.

| Parameters | Experimental Diets |  |  |  | TWO-WAY ANOVA p-value |  |  |
| --- | --- | --- | --- | --- | --- | --- | --- |
|  | D1 (SE) | D2 (SS) | D3 (FE) | D4 (FS) | Feed processing type | Protein source | Interaction |
| Lipid (%) | 5.58 ± 0.27<sup>ab</sup> | 6.36 ± 0.66<sup>bc</sup> | 5.33 ± 0.16<sup>a</sup> | 6.79 ± 0.61<sup>c</sup> | 0.029 | 0.803 | 0.370 |
| Crude Protein (%) | 67.28±1.98 | 66.19±0.64 | 69.08±4.10 | 66.71±2.98 | 0.305 | 0.483 | 0.698 |
| Ash (%) | 11.99±0.99 | 12.13±0.64 | 11.30±0.56 | 12.48±2.14 | 0.387 | 0.815 | 0.491 |

Values are means of triplicate groups and presented as mean ± SD. Values in the same row having different superscript letters are significantly different ( $p < 0.05$ ). The lack of superscript letter indicates no significant differences among treatments.

Shrimp amino acid analysis (Table S3- Supplementary Materials) revealed significant differences in lysine and isoleucine contents, particularly lower in D2 (SS), and lysine content was higher in D3 (FE) (p<0.05). Feed processing type significantly influenced valine content (p<0.05) with higher content noted in D1 (SE) and D3 (FE). Likewise, within each protein source, all shrimp fed extruded diets were significantly higher in isoleucine and lysine (p<0.05). Table S4 (Supplementary Materials) displayed the fatty acid contents within the sampled shrimp. D3 (FE) had significantly higher levels of C16, while D1 (SE) and D2 (SS) had higher n-6 polyunsaturated fatty acids (C18:2n-6, C18:3n-6, C20:2n-6, C20:3n-6) (p<0.05). Fishmeal–based diets D3 (FE) and D4 (FS) had higher n-3 polyunsaturated fatty acids content (C20:5n-3, C22:6n-3). No significant differences were observed in dimethylacetals among the diets (p>0.05).

#### 3.3. Feeding behaviour - Movement

Frame-by-frame imaging was assessed daily to quantify shrimp feeding behaviour, and dietary impacts on behaviour at salinities of 30 ppt and 5 ppt were analysed (Table 5). At 30 ppt, attraction to feed (Figure 3) was highest for diet D3 (FE), significantly outperforming D1 (SE) and D2 (SS) (p <0.05). This trend persisted at 5 ppt, with D3 (FE) and D4 (FS) showing higher attraction than D1 (SE) and D2 (SS) (p <0.05). A two-way ANOVA confirmed significant effects of protein source on feed attraction (p<0.05). Regarding feeding time, at 30 ppt, D3 (FE) induced longer durations compared to D1 (SE), D2 (SS) and D4 (FS) (p<0.05). At 5 ppt, D3 (FE) and D4 (FS) showed significantly higher feeding times compared to D1 (SE) and D2 (SS) (p <0.05). Feed processing type and protein source notably influenced feeding duration, with a preference for fishmeal–based feed (p <0.05). Shrimp on diets D1 (SE) and D2 (SS) displayed higher transition rates and were further from feed compared to those on D3 (FE) and D4 (FS), across both salinities (p <0.05). Significant impacts of feed processing and protein source were observed in transitions and distance-to-feed (p <0.05).

**Figure 3.**
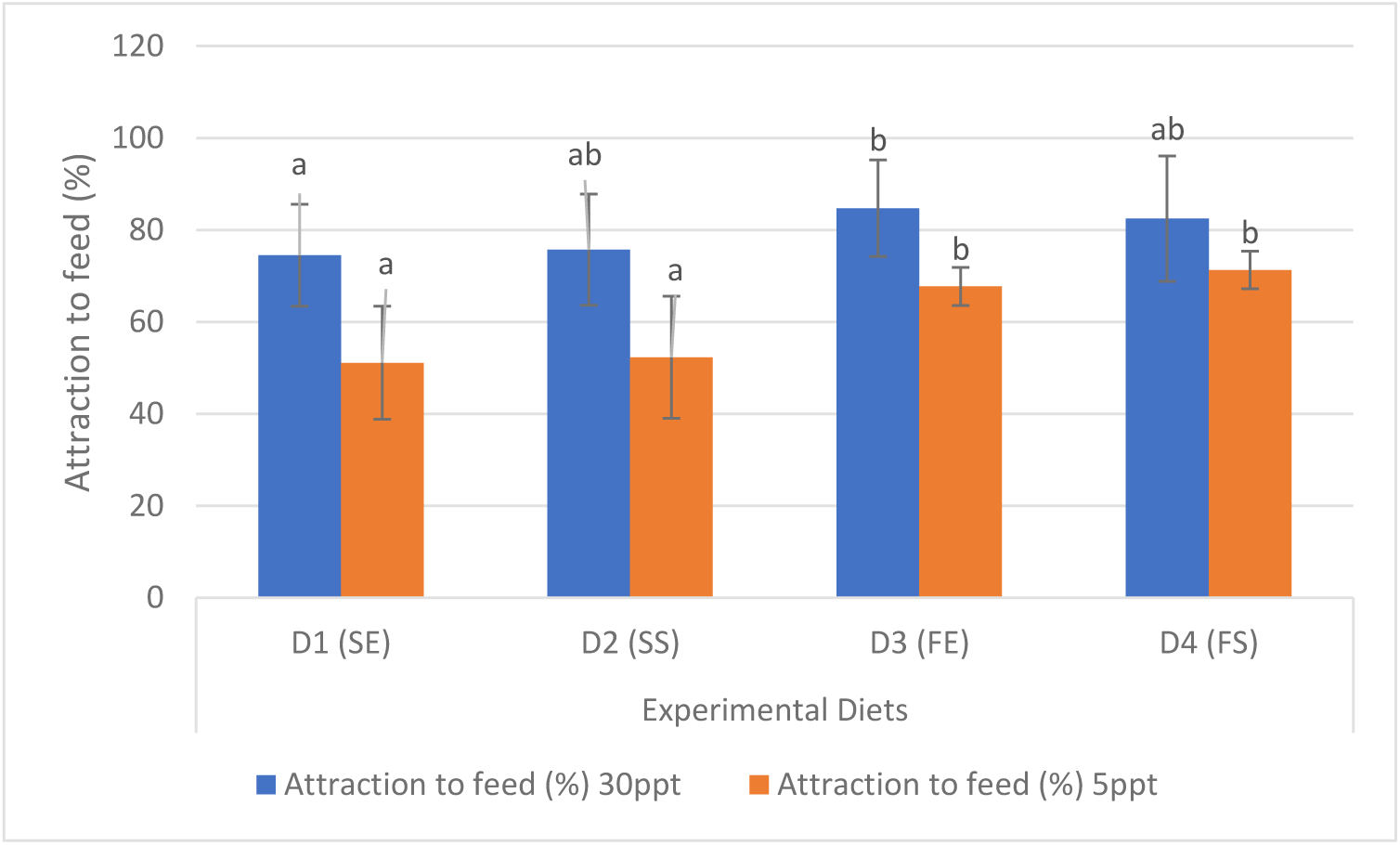
Attraction to feed at two different salinities. D1 (SE) = soya and pea extruded, D2 (SS) = soya and pea steamed, D3 (FE) = Fishmeal extruded and D4 (FS) = fishmeal steamed.

**Table 5.**
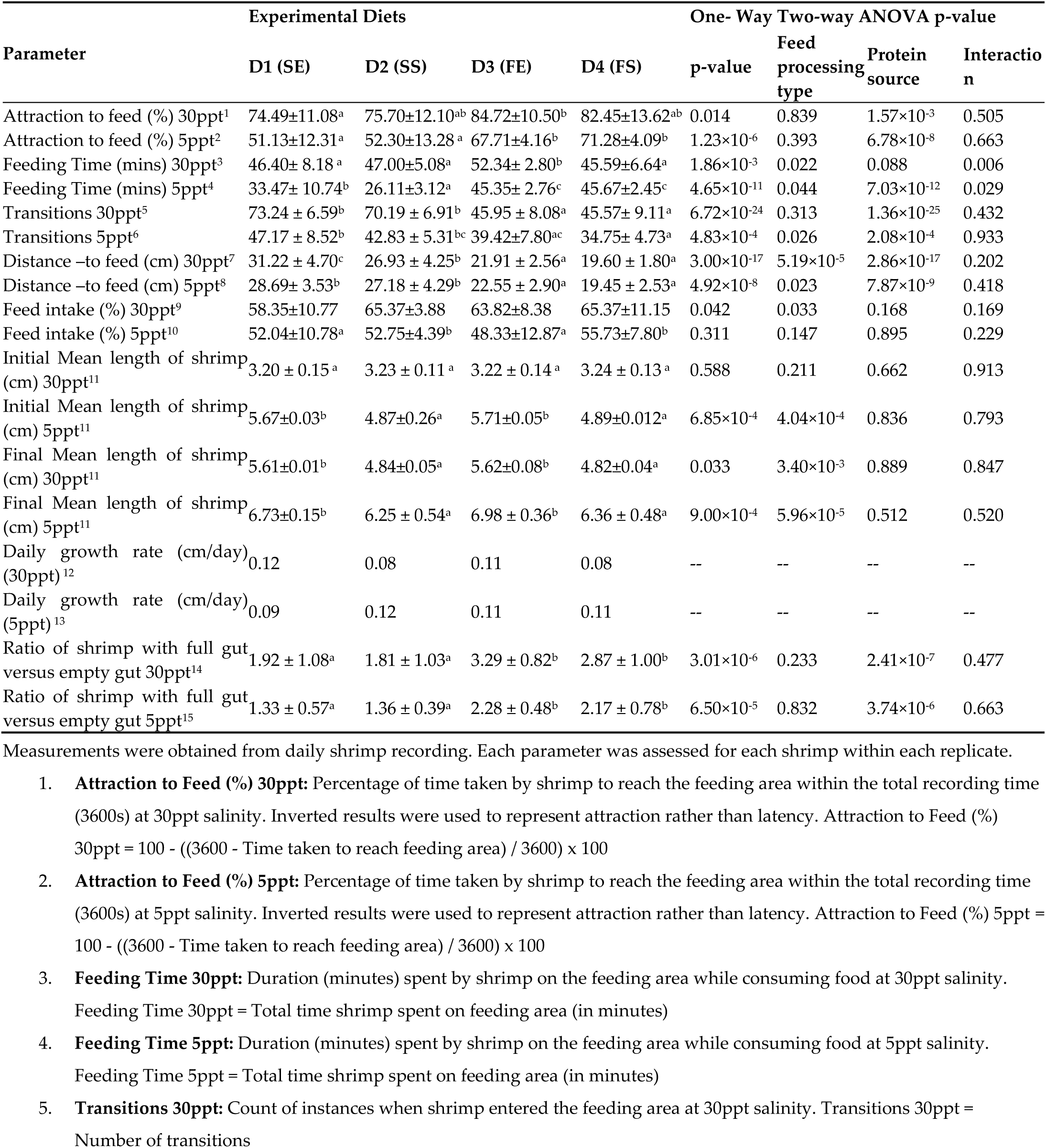
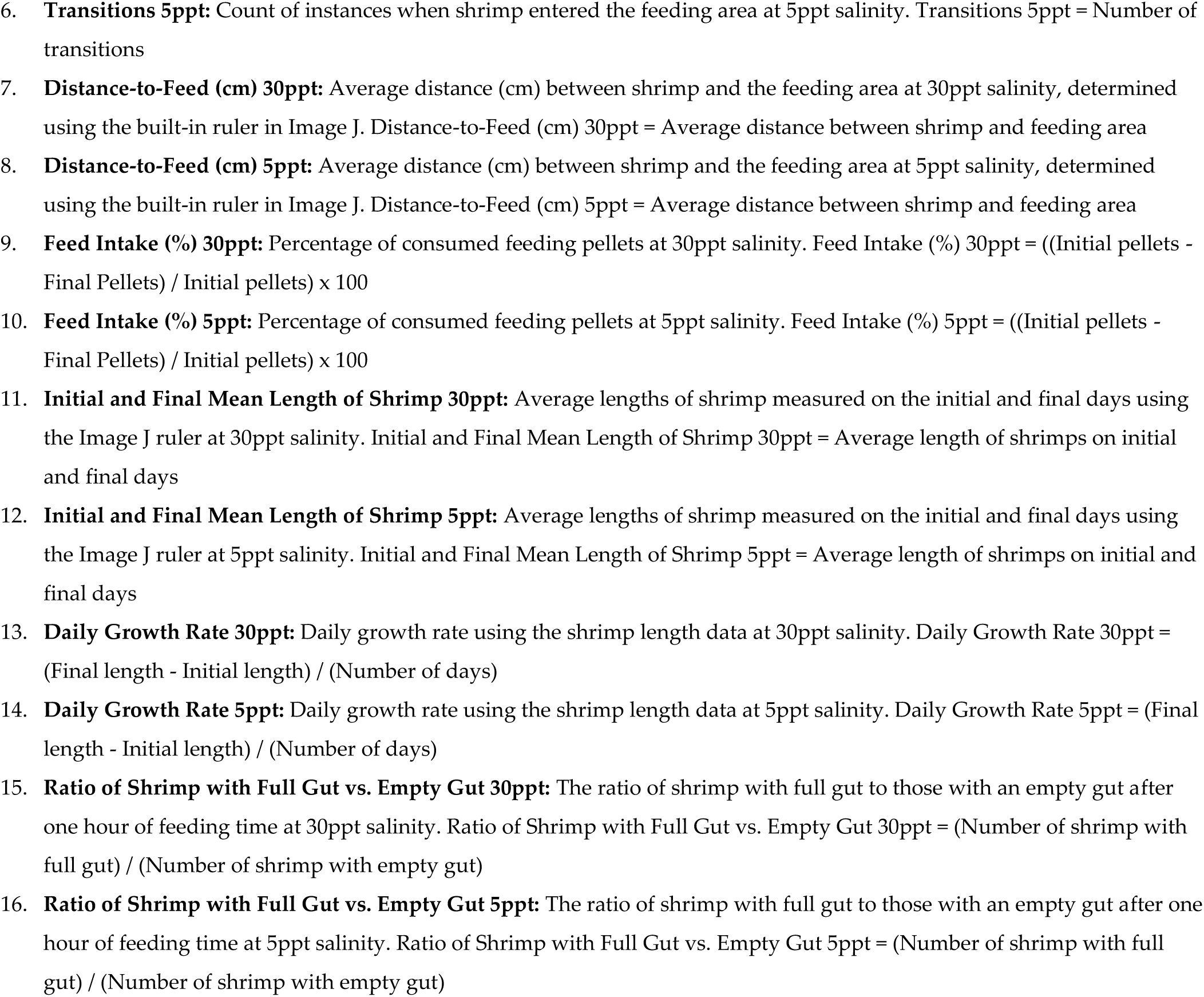
Image analysis parameters of *L. vannamei* fed the experimental diets for 33 days of the trial showing differences in behaviour at different salinities (30 ppt and 5 ppt). D1 (SE) = soya and pea extruded, D2 (SS) = soya and pea steamed, D3 (FE) = Fishmeal extruded and D4 (FS) = fishmeal steamed.

Feed intake varied with diet. At 30 ppt, D2 (SS) and D4 (FS) showed the highest intake, while it was lowest for D1 (SE), however, these values were not significant (p>0.05). At 5 ppt, D2 (SS) and D4 (FS) had significantly higher feed intake (p>0.05). Feed processing type alone significantly affected intake at 30 ppt (p <0.05). In terms of growth, shrimp fed D1 (SE) and D3 (FE) were longer than those on D2 (SS) and D4 (FS) at 30 ppt and 5 ppt (p <0.05). A two-way ANOVA highlighted significant effects of feed processing type on final shrimp length at both salinities (p <0.05). Daily Growth Rates varied with diet and salinity, but no consistent pattern was observed. Higher gut fullness ratios were seen in shrimp fed D3 (FE) and D4 (FS) at both salinities, indicating a preference for fishmeal–based feed (p <0.05).

#### 3.4. Feeding behaviour - Sound

The hydrophone data analysis revealed significant variations in clicking sound duration among shrimp diets (Table 6, Figure 4 and Figure 5). From a representative sample, shrimp fed D1 (SE) registered the most clicks (505), followed by D3 (FE) (380 clicks), D2 (SS) (265 clicks), and D4 (FS) (149 clicks), highlighting differences in click rates based on feed processing types. Shortest durations were observed in shrimp fed D1 (SE) (17.97 ± 11.67 ms) and D2 (SS) (17.24 ±10.73 ms), while the longest were in D3 (FE) and D4 (FS) (20.10 ± 13.04 and 23.89 ± 18.31 ms, respectively) with a small effect size (p < 0.05, *ω*^2^= 0.022). Low-frequency clicking sounds (<20 kHz) varied significantly, with D2 (SS) recording the highest frequency (15.13 ± 0.75 kHz) and D4 (FS) the lowest (14.59 ± 1.12 kHz), indicating a small effect size (p < 0.05, *ω*^2^= 0.022). In contrast, high-frequency sounds (>20 kHz) differed, with D1 (SE) having the lowest (24.97 ± 0.94 kHz) and D4 (FS) the highest frequencies (25.34 ± 1.40 kHz), showing a small effect size (p < 0.05, *ω*^2^= 0.012). Peak frequencies also significantly varied, with D4 (FS) having the lowest (20.14 ± 2.18 kHz) and D2 (SS) the highest (20.68 ± 1.37 kHz) peak frequencies (p < 0.05, *ω*^2^ = 0.009, very small effect size). However, energy (dB) and RMS values showed no significant differences across diets (p > 0.05).

**Figure 4.**
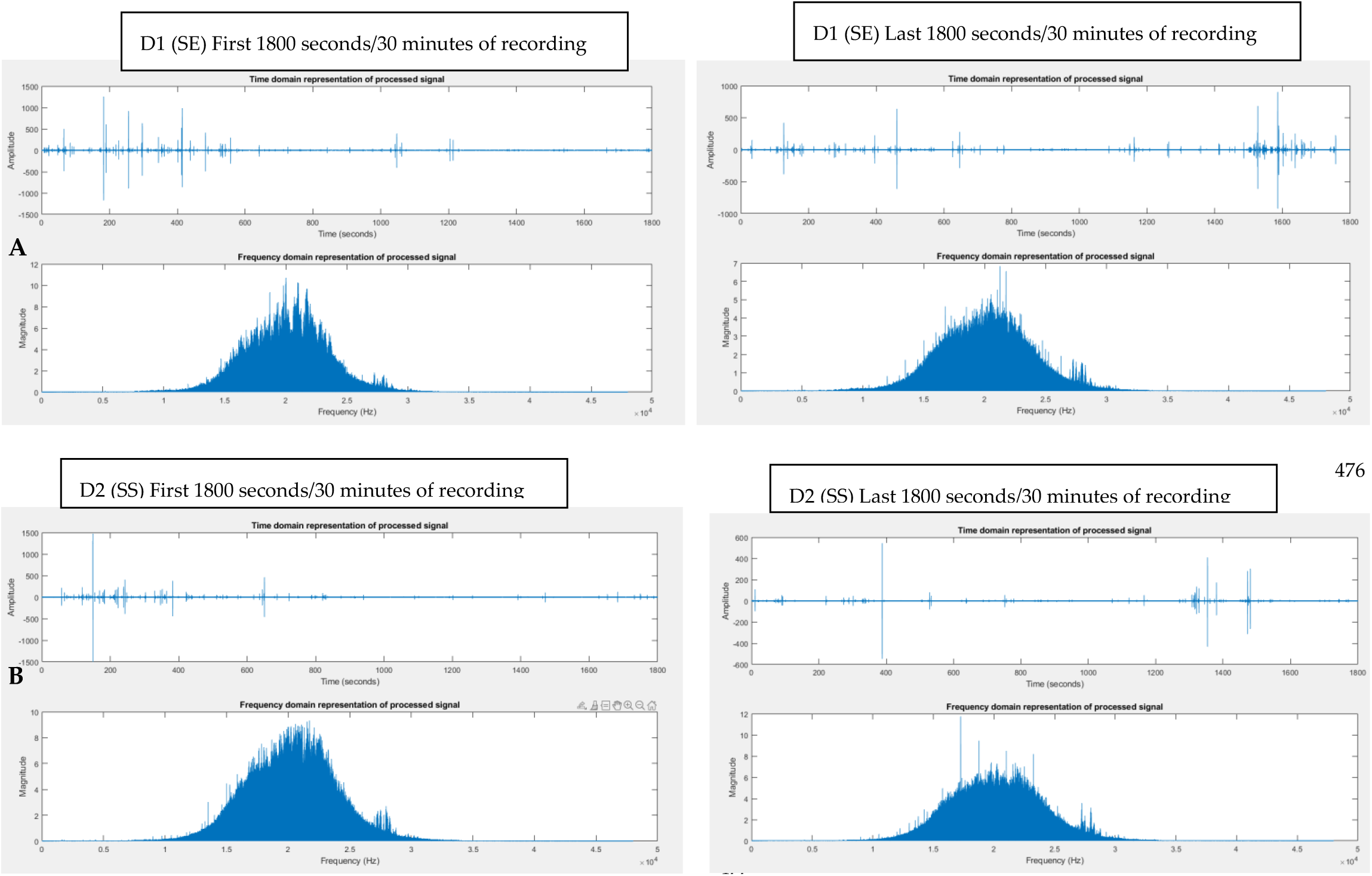
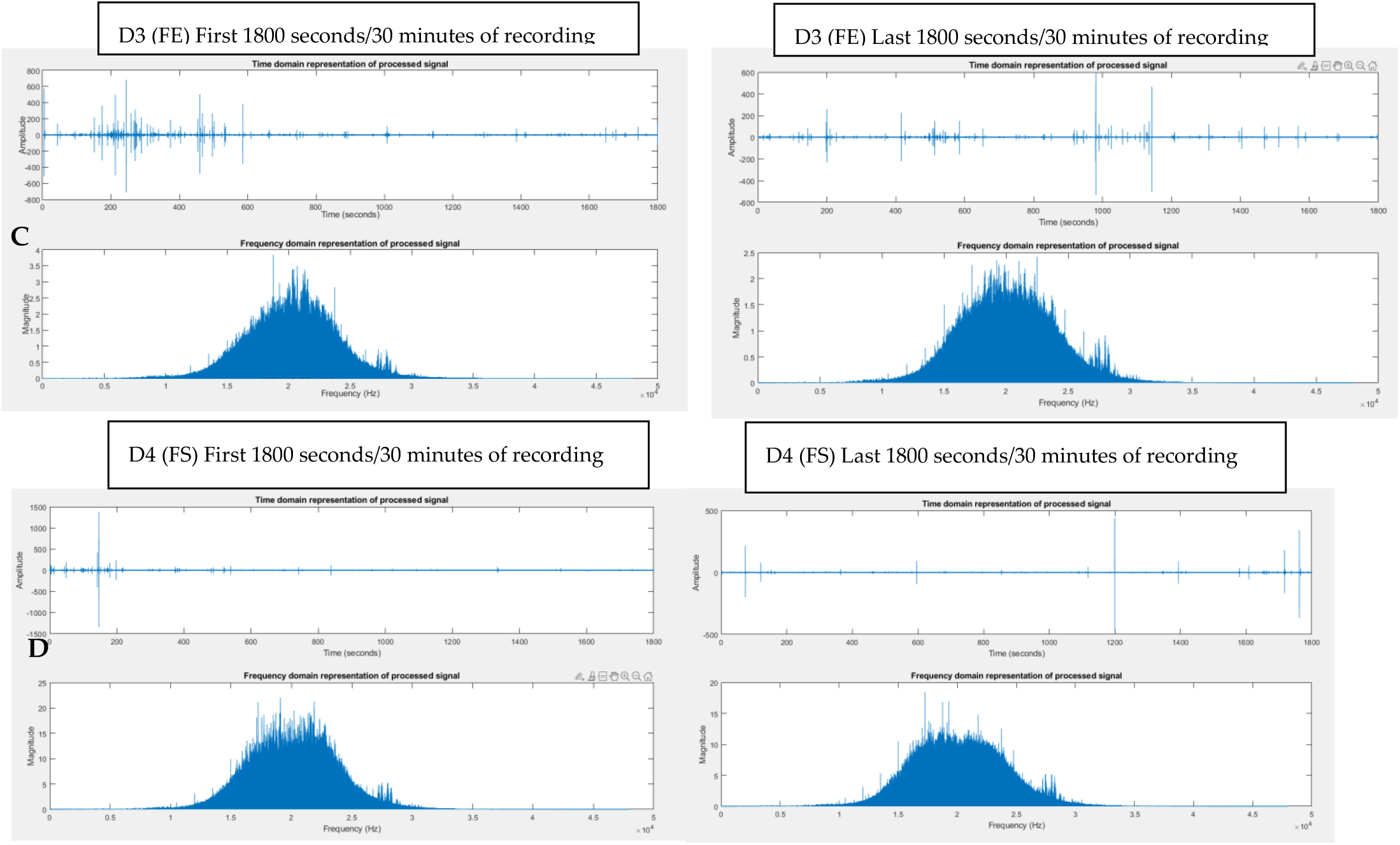
Example of Click Analysis and Filtration Results for D1 (SE) = soya and pea extruded (A), D2 (SS) = soya and pea steamed (B), D3 (FE) = Fishmeal extruded (C) and D4 (FS) = fishmeal steamed (D). The top graph shows the time domain representation of the acoustic signal. X-axis shows the time scale, whereas the signal’s amplitude is shown on y-axis. Large peaks in this time domain plot represent the clicking events. The bottom graph shows the frequency spectrum of the initial 1800 seconds (left image) and the final 1800 seconds (right image) of processed signal from examples of each dietary treatment group. These graphs show the pre-processed signal which is passed through the filter.

**Figure 5.**
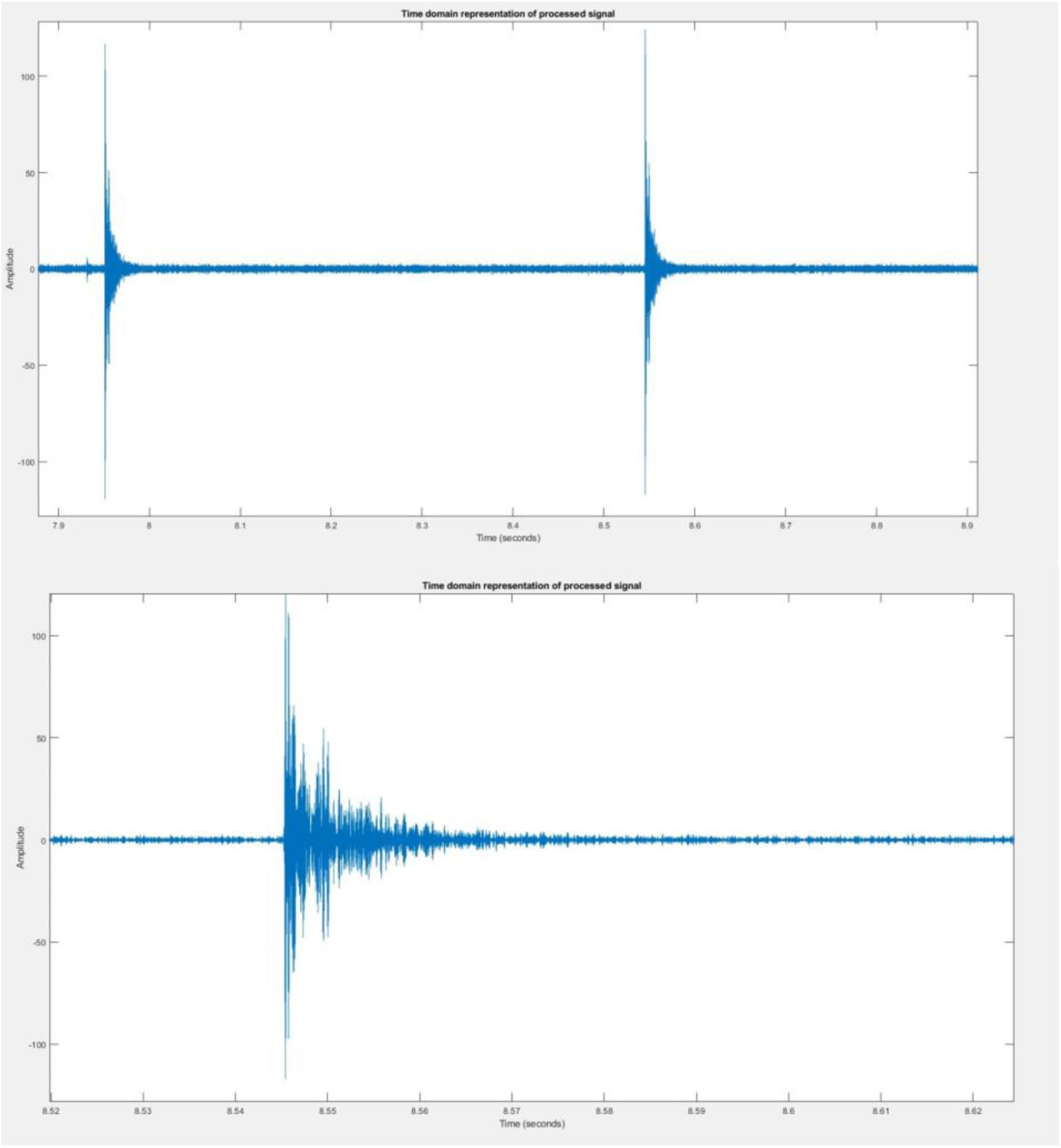
Zoomed examples of shrimp clicks.

**Table 6.**
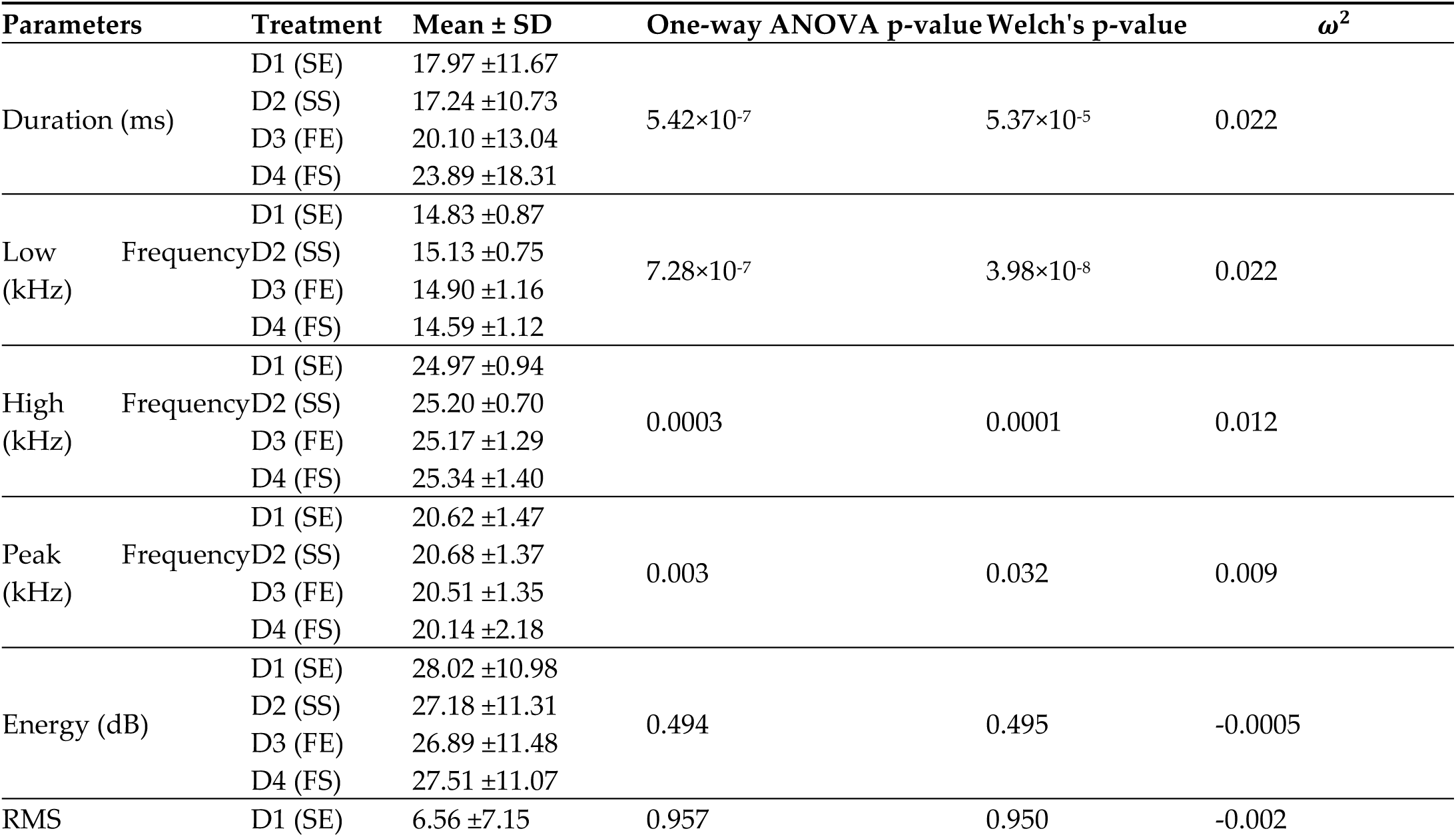
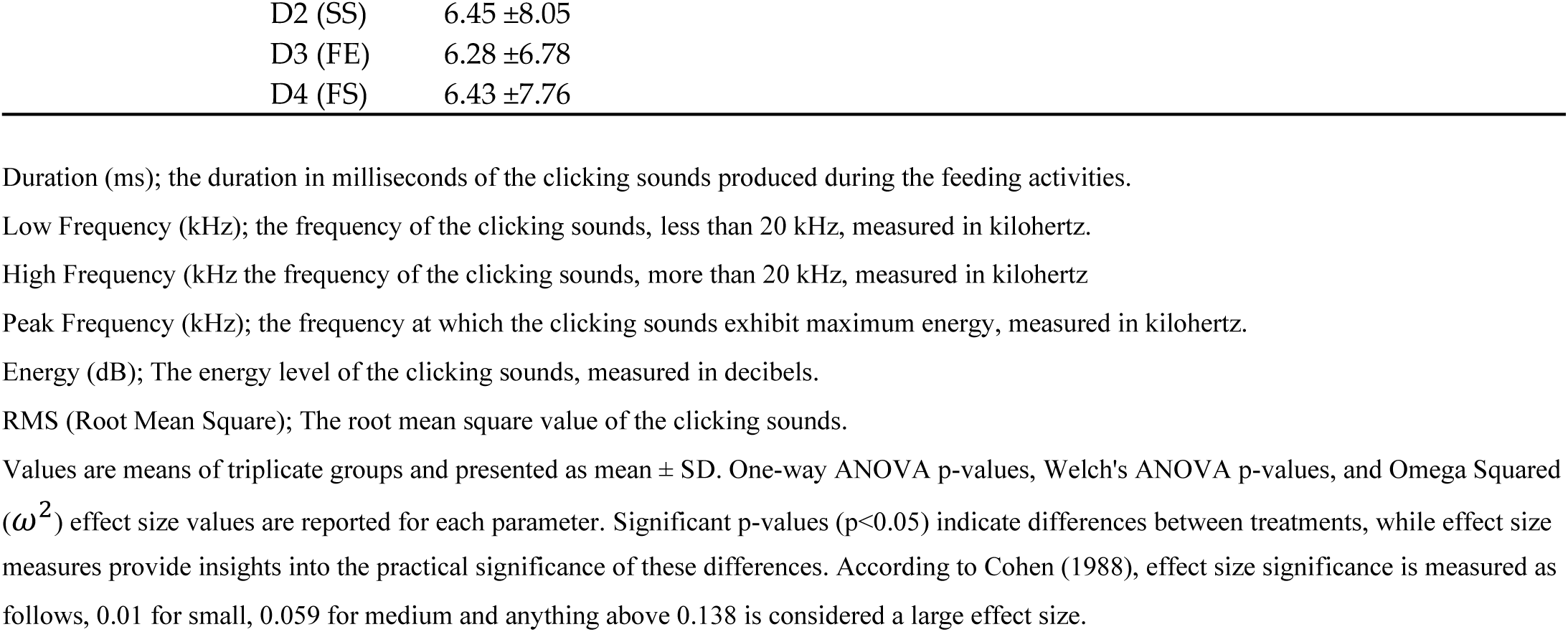
Click signal statistical analysis of *L. vannamei* fed the experimental diets D1 (soya and pea extruded), D2 (soya and pea steamed), D3 (Fishmeal extruded) and D4 (fishmeal steamed).

**Figure 6.**
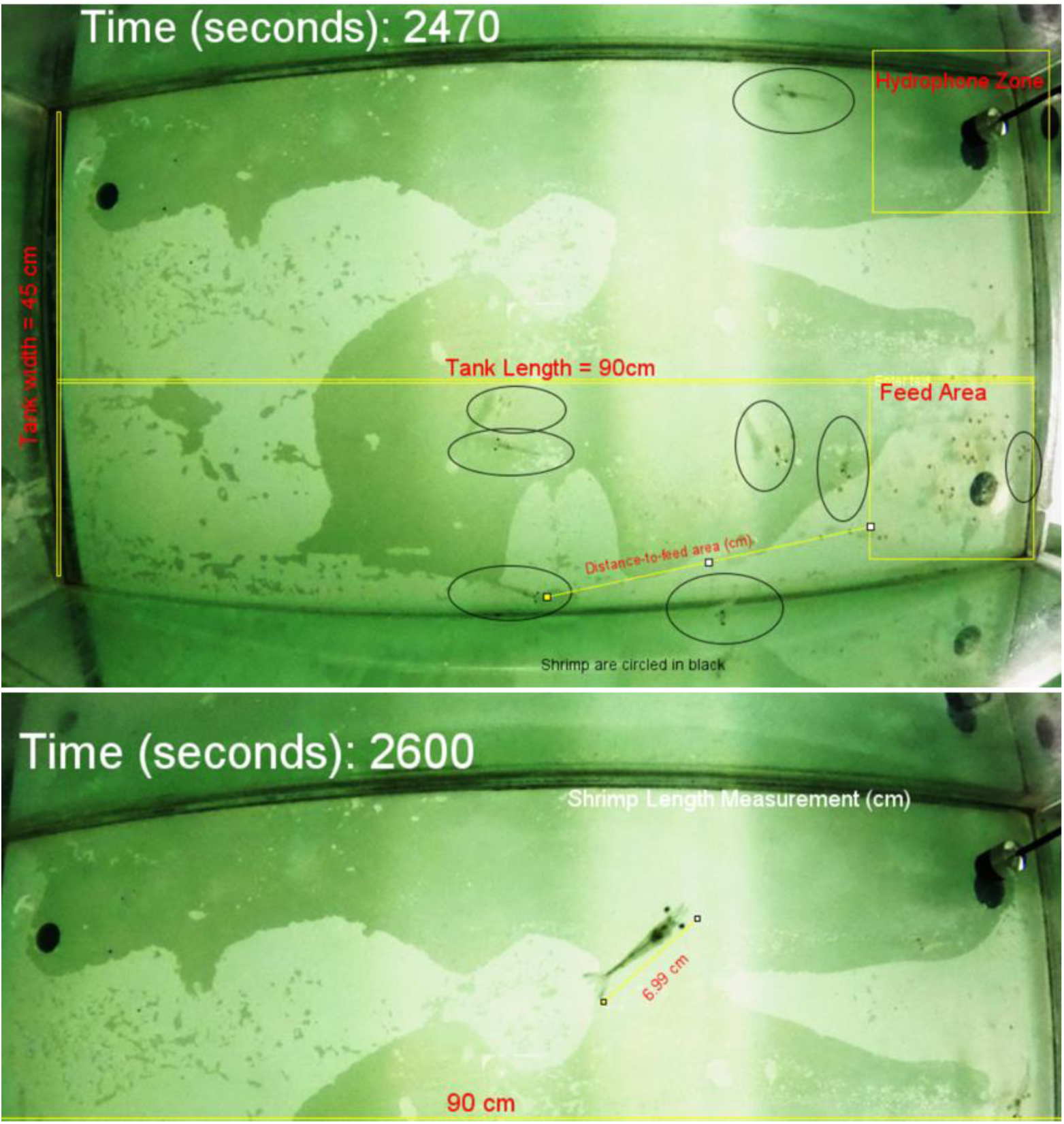
Image Frame on Image J showing the Distances and Shrimp Length Measured with shrimp circled in Black.

## 4. Discussion

This research advances the integration of imaging and acoustic techniques, following Darodes de Tailly et al. (2021), to look at the effects of protein source (soya and pea proteins vs. fish meal), processing types (steamed pellets vs. extruded pellets), and salinity (normal-30 ppt followed by stress- 5 ppt) on *L. vannamei* feeding behaviour, building on the foundation laid by earlier studies (Samocha et al. 2004; Smith et al. 2005). Unlike previous studies, this integration involving nutritional analyses, imaging and acoustic data, offers a more comprehensive understanding of shrimp responses, which is crucial for developing more effective and sustainable aquaculture practices.

### 4.1. Effects of feed processing type and protein source on shrimp diets

Although designed as isolipidic and isonitrogenous, the diets in this study exhibited nutritional variations. Soya/pea protein diets (D1 SE and D2 SS) had higher lipid content than fishmeal protein diets (D3 FE and D4 FS), with extruded feeds showing higher lipid content overall. This is consistent with findings that plant proteins like soybean enhance oil absorption in pellets, especially extruded ones (Samuelsen et al., 2013; Bektursunova, 2023; Xing et al., 2023). These variations can impact shrimp growth, by affecting lipid metabolism and energy utilization, which are vital for shrimp growth and immune response. Extrusion is known to boost digestibility and nutrient availability (Hilton, Cho, and Slinger, 1981; Vens-Cappell, 1984; Booth et al., 2002; Adedeji et al., 2015), while steaming increases moisture and pliability, reflected in the higher moisture content of steamed pellets (D2 SS and D4 FS), with the lowest moisture in fishmeal-based extruded feed (Diet FE).

Protein sources in the diets influenced their amino acid profiles. Fishmeal’s high-quality protein with balanced essential amino acids contributes to better growth and feed utilization in aquatic animals (Hua et al., 2019; Malcorps et al., 2019). Supplemented amino acids have also been studied for their ability to leach into the water, which can be a problem in shrimp feed due to their slow feeding behaviour (Prado et al., 2016). Despite being supplemented into the soya and pea–based diets, the levels of lysine, threonine and methionine were lower in Diets D1 (SE) and especially D2 (SS). Previous studies have shown that synthetic amino acids had higher leaching rates (Zarate and Lovell, 1997) with a study stating that 15% of this synthetic lysine was lost within 15 seconds of being immersed (Prado et al., 2016) and levels of free crystalline amino acids in feeds could be reduced by 80% within minutes of being exposed to water (Lopez-Alvarado et al., 1994). The amino acid profiles, particularly the lower levels of lysine, threonine, and methionine in plant-based diets, despite being supplemented, have implications for shrimp growth and survival. The leaching rates of synthetic amino acids, as highlighted by Zarate and Lovell (1997), suggest a potential reduction in the nutritional value of feeds in water, which could lead to suboptimal growth rates and increased susceptibility to diseases. Future studies should explore how these rates impact feed detection and their overall ecological implications.

Fatty acid composition also varied among the diets. Plant–based diets (D1 and D2) had lower proportions of SFAs and MUFAs but higher n-6 PUFAs, typical of plant oils like soybean. Conversely, fishmeal–based diets (D3 and D4) had higher n-3 PUFAs, reflecting fishmeal inclusion. These findings align with Tocher et al. (2019) and Rizzo et al. (2023). A comparative analysis or meta-analysis with similar studies could provide insights into optimizing diet formulations for enhanced shrimp growth and environmental benefits. Diets rich in n-3 PUFAs (D3 and D4) can enhance the immune system and improve stress tolerance in shrimp, while higher n-6 PUFA levels (D1 and D2) might lead to an imbalance in fatty acid ratios, potentially affecting inflammatory responses and overall health. Additionally, variations in TBARS levels between diets could indicate differences in diet oxidative stability. Higher TBARS levels in fishmeal–based diets suggest greater oxidation susceptibility of n-3 PUFAs (Van Hecke et al., 2021).

### 4.2. Effects of feed processing type, protein source and salinity on shrimp growth and body composition

In this study, contrasting results were found in the impacts of extrusion and steam- pelleting on aquatic organisms, echoing mixed findings in previous research. Hilton et al. (1981), Vens-Cappell (1984), and Adedeji et al. (2017) noted improved growth in species like rainbow trout and *L. vannamei* when fed extruded pellets. Conversely, Booth et al. (2002) observed better outcomes with steam-pelleted diets in silver perch. The current study showed that while extruded feeds (D1 SE and D3 FE) led to numerically higher final body weights and significantly greater mean lengths, steam-pelleted diets (D2 SS and D4 FS) resulted in higher survival rates, particularly at lower salinities.

Low salinity challenged the shrimp’s physiological functions, necessitating adjustments in internal osmotic balance (Charmantier et al., 1991; Péqueux, 1995; Pham et al., 2012). The contrast in shrimp survival rates between extruded and steam-pelleted diets under different salinity conditions suggests that feed processing affects shrimp’s ability to cope with environmental stress (Figure 1). This would be particularly evident in instances of intense rain and flooding when salinity levels are known to dramatically decrease. The higher mortality in shrimp fed with extruded pellets at low salinity could be attributed to the increased energy expenditure for osmoregulation, leaving less energy for feeding and growth. If extruded pellets require more energy to consume due to their harder texture, shrimp are less likely to ingest them in lower salinities due to the increased energy required for osmoregulation, which could lead to a decrease in the uptake of the necessary nutrients to support proper osmoregulation and stress responses resulting in increased mortality. Camacho-Jiménez et al. (2018) highlighted the role of crustacean hyperglycemic hormones (CHHs) in osmo-ionic regulation under acute salinity stress, suggesting the increased energy requirement for osmoregulation could reduce nutrient uptake in shrimp, impacting their survival. Despite the higher mortality in low salinity conditions, shrimp on extruded diets exhibited higher growth parameters, likely due to minimal amino acid leaching compared to steam-pelleted diets. The protein source in the diets significantly influenced shrimp amino acid profiles. Fishmeal is known for its balanced amino acids, contributing to better growth (Wouters et al., 2001; Glencross et al., 2002; Gonzalez-Galaviz et al., 2020). However, the plant-based diets in this study showed higher rates of amino acid leaching (Prado et al., 2016), possibly affecting shrimp feed efficacy.

Furthermore, different protein sources in the diets led to varied fatty acid profiles in the shrimp. The use of fishmeal in diet D3 (FE) compared to plant–based proteins in diets D1 (SE) and D2 (SS) likely resulted in a higher level of C16:0 (palmitic acid) due to the naturally higher content of this fatty acid in fishmeal. This difference in dietary fatty acid composition influences the fatty acid synthesis and accumulation in shrimp tissues. Past research suggests that extrusion processing could modify the lipid structure of pellets, influencing fatty acid composition (Imran et al., 2015), and this is supported by the observed differences between D1 (SE) and D3 (FE). D3 (FE) and D4 (FS) had higher levels of MUFAs (C16:1n-7, C18:1n-7, C20:1n-11, and C20:1n-9), possibly due to the use of fishmeal protein.

Additionally, previous findings show that dietary lipid levels are crucial for growth in low-salinity environments (Jannathulla et al., 2019; Chuphal et al., 2021). The current study showed that diets made through extrusion have higher lipid and energy content. Higher dietary lipid levels have been previously correlated with increased growth in lower salinity levels (Jannathulla et al., 2019; Chuphal et al., 2021; Jana et al., 2022). Likewise, a study by Chen et al. (2015) revealed significant changes in gene expression in the hepatopancreas of *L. vannamei* exposed to long-term low salinity with key affected pathways in lipid metabolism and osmoregulation. Sui et al. (2015) and Jannathulla et al. (2019) found that optimal dietary protein levels and lipid contents vary with salinity, and different lipid/essential fatty acid (EFA) levels in the diet influenced the haemolymph protein levels of *L. vannamei*, indicating a potential alteration in protein synthesis and enzyme activity related to lipid metabolism under varying salinity conditions.

Huang et al. (2019) and Chen et al. (2019) revealed that lipidomic responses and fatty acid metabolism in *L. vannamei* are significantly influenced by environmental salinity and dietary fatty acid sources, affecting growth and survival. Likewise, Chuphal et al. (2021) found that higher dietary lipid levels in *L. vannamei* diets led to increased lipase activity, indicating a direct influence on lipid metabolism enzymes. The current study also highlighted how protein source and feed preparation method influence shrimp lipid content, with fishmeal steamed diets leading to the highest lipid accumulation in shrimp. Shrimp’s fatty acid composition is vital for their osmoregulatory capacities and has implications for human health and the environment. Rich in beneficial LC-PUFAs like EPA and DHA, shrimp can offer cardiovascular and cognitive benefits to human consumers (Swanson et al., 2012; Fang et al., 2022). Diet manipulation can enhance shrimp’s nutritional value. However, fishmeal sustainability concerns, due to overfishing and ecosystem impacts, have prompted the exploration of more sustainable protein sources like soy and pea proteins. The choice of protein source, as shown in this study, affects shrimp’s fatty acid composition, thus influencing the nutritional quality of farmed shrimp for human consumption. The current study’s insights into the effects of feed processing and salinity on shrimp growth highlight the need for tailored feed strategies in shrimp farming. These strategies should aim to maximize growth and survival rates, particularly under varying environmental conditions.

### 4.3. Effects of feed processing type, protein source and salinity on shrimp feeding behaviour

Salinity is known to significantly affect the physiology and behaviour of aquatic organisms, including shrimp (Kefford et al., 2012; Bal et al., 2022). Duan et al. (2022) found that salinity and dissolved oxygen (DO) concentrations significantly affect the tail-flip speed (a measure of swimming ability) of *L. vannamei*. They observed that the tail-flip speed increased and then decreased with increasing salinity, indicating an optimal salinity range for maximum swimming ability. Additionally, different salinities and DO concentrations directly affected shrimp physiology, inducing changes in metabolite concentrations in haemolymph, hepatopancreas, and abdominal muscles, which are linked to energy consumption and exercise ability. In the current study, two salinity levels (30 ppt and 5 ppt) elicited significant differences in the parameters, ‘attraction to feed’ and ‘feeding times’. Shrimp showed increased attraction and prolonged feeding times at 30 ppt, which has been correlated with enhanced osmoregulation and metabolic efficiency, typically observed at higher salinities (Charmantier et al., 1989; Lignot et al., 2000; Bückle et al., 2006). This suggests that shrimp can optimize nutrient intake and utilization under these conditions. This behavioural adaptation likely contributes to improved metabolic efficiency, essential for growth and survival in fluctuating environmental conditions (Péqueux, 1995). However, this behaviour may change during the shrimp’s life cycle (Charmantier, 1991). The study’s findings are in agreement with research on decapods with biphasic life cycles, where morphological and eco-ethological shifts during ontogenetic transitions are crucial for adaptation to specific environments (Charmantier et al., 1991; Anger, 2006; Pham et al., 2012). These transitions, particularly under osmotic stress, affect survival rates due to energy allocation for osmoregulation, potentially influencing feeding behaviour (Rahi et al., 2021).

The type of feed and protein source also influence nutrient availability and palatability, thereby affecting feeding behaviour in aquatic species (Hua et al., 2019). In the current study, variations in feeding behaviour were observed in response to different feed processing types (extruded and steamed pellets) and protein sources (soya/pea meal and fishmeal). Shrimp fed extruded pellets demonstrated enhanced attraction to feed, likely due to increased palatability and nutrient availability (Tantikitti, 2014). Shrimp possess several specialized sensory structures including chemoreceptors, which play a crucial role in detecting chemical cues in their environment, such as from food (Dall et al., 1990; Alday-Sanz, 2010; Schmidt & Mellon, 2011; Kamio & Derby, 2017; Nguyen et al., 2018). These chemoreceptors are found on the setae and bristles on their mouthparts and other appendages. Notably, chemoreceptors located on the lateral flagellum of the antennules can detect dissolved compounds released from the feed, such as amino acids, peptides, and other organic molecules (Dall et al., 1990; Gadient and Schai, 1994; Obaldo et al., 2006; Alday-Sanz, 2010; Eap et al., 2020). Shrimp fed fishmeal–based diets exhibited longer feeding times and higher attraction to feed, potentially due to fishmeal’s amino acid profile (Browdy et al., 2006). The amino acid composition of diets has been shown to impact feeding behaviour and feed intake in aquatic species (Alam et al., 2004; Simon et al., 2021).

The differences in feeding behaviour correlated with growth performance indicated that favourable feeding environments characterized by higher salinities and nutrient-rich diets lead to better growth and higher biomass. Considering the trend towards reducing reliance on fishmeal in aquafeeds, it is essential to recognize how substituting fishmeal with other ingredients affects feeding behaviour and nutrient utilization. Feed consumption differed slightly between diets, suggesting the higher palatability of fishmeal–based diets. As shown in the current study, and previous studies, shrimp fed extruded pellets had higher final body weights and significantly higher body lengths despite lower feed intake, attributed to better water stability and nutrient retention in extruded pellets (Hoyos et al., 2017; Jescovitch et al., 2018; Soares et al., 2021). The slow feeding behaviour of shrimp can result in increased nutrient leaching and loss prior to ingestion, a crucial consideration when substituting fishmeal with alternative components like soybean, which may require the supplementation of essential crystalline amino acids (CAAs), such as lysine (Simon et al., 2021). The observed variations in feeding behaviour have significant implications for shrimp physiology. It is crucial to link these behavioural changes to the growth and metabolic efficiency of shrimp, potentially leading to improved feeding strategies in aquaculture.

Peak feeding activity, characterized by more frequent clicks, was observed within the first 10 minutes of feed being dispensed, after which the number of clicks reduced, which is in line with observations by Hamilton et al. (2023). Clicking sounds may be influenced by the texture and hardness of feed particles, as well as shrimp size in the case of the snapping shrimp, with larger shrimp correlated with louder snaps due to larger frontal claws as they age, increasing in sound pressure levels and energy (Au et al., 1998). Click signal analysis provided insights into the acoustic behaviour of *L. vannamei* in response to different feeds. Differences in the number of clicks and click duration suggest variations in feeding efficiency and consumption rates between diets. The findings show that the clicking sounds were between 15 kHz and 25 kHz and they varied significantly between the diets with Diet 1 (SE) having the shortest duration and Diets 3 (FE) and 4 (FS) having the longest durations. The physical characteristics of the feed, such as pellet hardness and density, impact the clicking behaviour of *L. vannamei* (Peixoto et al. 2020). The click duration recorded in the current study (14.0 -15.0 ms) is almost three times the values previously reported (4.7 ms) by Silva et al. (2019). This difference is probably related to the highly reflective glass tank surface (90 cm × 45 cm X 20 cm) used in the current study, in contrast to smaller tank surface used by Silva et al. (2019) and this could result in an exaggeration of the clicking sound duration owing to sound reverberation (Bart et al., 2001). Also, Silva et al. (2019) used 6 shrimp per tank which is a smaller number compared to 10 shrimp per tank used in this current study. Therefore, the difference in click duration might also be related to number of shrimp per tank, causing overlapping signals. There are other parameters of acoustic monitoring that are useful for assessing shrimp feeding behaviour, such as measuring the click frequencies.

The differences in low, high, and peak frequencies suggest different feeding patterns with different diets. For instance, shrimp fed D2 (SS) exhibited the highest low frequency sounds and shrimp fed D4 (FS) exhibited the highest high frequency sounds suggesting variations in shrimp behaviour and feeding efficiency based on feed processing type and protein source. The peak frequency of the clicks emitted by shrimp offered extruded diets in this research is notably lower than those fed steamed diets. This disparity could be linked to the food texture, as mechanically robust feeds tend to exhibit lower peak frequencies, as previously observed (Duizer, 2004). During the extrusion process, feed mixtures are subjected to high heat, pressure, moisture (110 –200°C, 350 –l400 psi, 20% - 30%, respectively) (Frame, 1994), and subsequently forcing the mixture through a narrow opening (die) to create a specific shape. The high temperature and pressures used in extrusion cooking contribute to gelatinization of starches and formation of dense and durable pellets which have a crunchy, hard texture. On the other hand, steam pelleting involves conditioning feed mixtures with steam involving moderate temperature, pressure, and moisture (< 90°C, 75 to 125 psi, 17% –18%, respectively) (Hilton et al., 1981; Booth, 2002; Adedeji, 2015) and then compressing through a pellet mill to form pellets with softer textures. In the current study, the clicking patterns of *L. vannamei* consistently exhibited a high peak frequency, surpassing 15 kHz across all diet combinations. These pronounced clicks could be advantageous for detecting feeding activity in aquaculture systems and encouraging rapid detection of feed. Hence, the focus should extend beyond optimizing systems geared to minimize acoustic disturbances in aquaculture (Bart et al., 2001; Radford and Slater, 2019) and consider enhancing the suitability of sonic efficient feeds in shrimp farms where acoustics are used to monitor feeding patterns. The study’s acoustic monitoring data, revealing longer and more frequent clicking sounds for fishmeal-based diets, suggests that the feed’s texture and composition influence shrimp feeding behaviour. This finding has implications for feed design, indicating a need for feeds that facilitate efficient nutrient intake and minimize energy expenditure during feeding.

Shrimp feeding behaviour is influenced by dietary composition, including feed processing type and protein source (Sanchez et al., 2005). Extruded feeds have different physical characteristics and nutrient availability compared to steamed pellets, affecting shrimp’s interaction with their food. Shrimp’s specialized mouthparts, adapted for different functions during feeding (Nguyen et al., 2018), respond differently to the texture of the feed, influenced by feed processing type and protein source. The texture of the feed, along with nutrient leaching and chemosensory responses, creates intricate interactions affecting shrimp feeding preferences and rates (Prado et al., 2016). Understanding these dietary factors’ impact on both the physical and chemical aspects of feed is crucial for optimizing feeding strategies in aquaculture. However, there are limitations in using imaging and acoustic techniques for assessing shrimp feeding behaviour, particularly in commercial-scale brown and green murky water and farm noise interference (Napaumpaiporn et al. 2013; Ullman et al. 2019). Future studies should aim to adapt these techniques for broader use in varied aquaculture environments. The simultaneous use of images and acoustics provides a comprehensive understanding of shrimp feeding behaviour, beneficial for aquaculture research during new feed trials.

## 5. Conclusions

This study highlights the profound impact of feed type on the behaviour and adaptability of *Litopenaeus vannamei* in two different salinity conditions. The findings reveal that fishmeal-based diets promote longer feeding times and greater attraction, likely due to their balanced amino acid profile and fatty acid composition, which are essential for shrimp growth and stress tolerance, further enhanced by better nutritional uptake, even under low-salinity stress. Additionally, acoustic data revealed that shrimp on fishmeal-based diets produced distinctly longer clicks whereas, extruded feeds led to more frequent clicking sounds, suggesting a strong link between diet and communication or stress behaviours. These findings emphasize the importance of tailored feed strategies that consider nutritional content, feed physical properties and environmental factors for optimal shrimp feeding. In practical terms, these insights can guide the development of more effective and sustainable shrimp feeds, balancing the nutritional needs of shrimp with environmental adaptability. For instance, optimizing the lipid content and amino acid profile in plant-based diets could enhance shrimp growth while reducing reliance on fishmeal, addressing sustainability concerns, but also increase feed attraction, thus reducing environmental impact. Moreover, understanding how feed processing affects feed palatability and shrimp energy expenditure can lead to the production of feeds that improve shrimp growth efficiency, particularly in challenging environmental conditions like low salinity. Future research should focus on exploring alternative sustainable protein sources, assessing their impact on shrimp feeding behaviour and the implications for environmental sustainability. Additionally, studies examining the long-term effects of varying feed types on shrimp behaviour and physiology would be valuable.

## Authors statement

AM: acquired funding, designed the experiment, drafted the manuscript, and analysed the data; SO: analysed feed behaviour using ImageJ; AB: acquired funding, contributed to experimental design, set up the experiment, developed image analysis methods in R and revised the manuscript; MN: formulated the feed, conducted the experiment, and revised the manuscript; SD: conducted the experiment; JS: formulated the feed, conducted the experiment.

## Data availability

Data will be made available upon request.

## Institutional Review Board Statement

The animal experiments in this study were conducted in accordance with the ethical guidelines and approved by the ethics committee of the University of Stirling (AWERB 2022 10523 7859).

## Acknowledgments

This work was supported by the Biotechnology and Biological Sciences Research Council [grant number BB/X511985/1]. The authors would like to sincerely thank Fiona Strachan, Charlotte Horrell, Kathryn Grace, Miriam Molina Moreno, Angus Beaton, and Chelsea Broughton who completed the nutritional analytical work at the Nutrition Analytical Service, Institute of Aquaculture, University of Stirling. The authors would also like to thank Dr Darren Green for assisting with the manuscript review.

