## Supplementary Tables for "Effects of Feed Processing Type, Protein Source, and Environmental Salinity on *Litopenaeus vannamei* Feeding Behaviour"

**Supplemental Materials**

Table S1. Amino acid profiles of experimental diets D1 (soya and pea extruded), D2 (soya and pea steamed), D3 (Fishmeal extruded) and D4 (fishmeal steamed). (g/100g)

| | **D1 (SE)** | **D2 (SS)** | **D3 (FE)** | **D4 (FS)** |
|---|---|---|---|---|
| **Aspartic acid** | 3.13 | 2.76 | 2.92 | 2.63 |
| **Serine** | 1.68 | 1.61 | 1.62 | 1.61 |
| **Glutamic acid** | 8.49 | 8.32 | 8.80 | 7.86 |
| **Proline** | 2.58 | 2.52 | 2.88 | 2.49 |
| **Glycine** | 1.53 | 1.47 | 2.36 | 2.08 |
| **Alanine** | 1.42 | 1.39 | 2.06 | 1.78 |
| **Valine** | 1.74 | 1.68 | 1.81 | 1.60 |
| **Isoleucine** | 1.68 | 1.61 | 1.71 | 1.52 |
| **Leucine** | 2.70 | 2.55 | 2.83 | 2.51 |
| **Tyrosine** | 1.23 | 1.16 | 1.27 | 1.13 |
| **Phenylalanine** | 1.86 | 1.71 | 1.71 | 1.54 |
| **Histidine** | 0.86 | 0.78 | 0.80 | 0.70 |
| **Lysine** | 2.10 | 1.94 | 2.43 | 2.19 |
| **Arginine** | 2.34 | 2.16 | 2.14 | 1.91 |
| **Threonine** | 1.14 | 0.73 | 1.29 | 1.15 |
| **Methionine** | 0.95 | 0.90 | 1.13 | 1.00 |
| **Cysteine** | 0.28 | 0.26 | 0.27 | 0.27 |

Table S2. Fatty acid profiles of experimental diets D1 (soya and pea extruded), D2 (soya and pea steamed), D3 (Fishmeal extruded) and D4 (fishmeal steamed). (% total fatty acids and mg FA.100$^{g-1}$)

| Fatty acids | **D1 (SE)** | | **D2 (SS)** | | **D3 (FE)** | | **D4 (FS)** | |
|---|---|---|---|---|---|---|---|---|
| | **%** | **mg. 100g$^{-1}$** | **%** | **mg. 100g$^{-1}$** | **%** | **mg. 100g$^{-1}$** | **%** | **mg. 100g$^{-1}$** |
| C14:0 | 1.86 | 88.71 | 1.83 | 72.64 | 2.12 | 84.43 | 2.01 | 80.04 |
| C15:0 | 0.37 | 17.56 | 0.35 | 13.78 | 0.42 | 16.92 | 0.42 | 16.74 |
| C16:0 | 13.70 | 652.28 | 14.01 | 557.26 | 16.29 | 649.66 | 16.93 | 674.91 |
| C18:0 | 2.47 | 117.49 | 2.63 | 104.52 | 2.20 | 87.91 | 2.40 | 95.63 |
| C20:0 | 0.32 | 15.15 | 0.36 | 14.32 | 0.19 | 7.69 | 0.22 | 8.71 |
| C22:0 | 0.20 | 9.43 | 0.27 | 10.93 | 0.23 | 9.33 | 0.23 | 9.19 |
| C24:0 | 0.07 | 3.53 | 0.10 | 4.11 | 0.13 | 5.08 | 0.15 | 5.86 |
| **ΣSFAs** | **18.99** | **904.15** | **19.54** | **777.56** | **21.59** | **861.03** | **22.36** | **891.09** |
| C16:1n-9 | 0.21 | 10.10 | 0.20 | 8.13 | 0.19 | 7.69 | 0.19 | 7.40 |
| C16:1n-7 | 2.19 | 104.42 | 2.11 | 83.86 | 3.27 | 130.61 | 3.23 | 128.87 |
| C18:1n-9 | 21.76 | 1036.07 | 21.85 | 869.50 | 17.15 | 683.90 | 17.14 | 682.99 |
| C18:1n-7 | 1.97 | 93.59 | 1.91 | 76.13 | 2.90 | 115.86 | 2.95 | 117.70 |
| C20:1n-11 | 0.28 | 13.52 | 0.28 | 11.12 | 0.70 | 28.09 | 0.74 | 29.57 |
| C20:1n-9 | 3.49 | 165.92 | 3.43 | 136.53 | 4.91 | 195.79 | 5.09 | 202.82 |
| C20:1n-7 | 0.22 | 10.32 | 0.19 | 7.74 | 0.30 | 12.14 | 0.32 | 12.68 |
| C22:1n-11 | 3.80 | 181.12 | 3.79 | 150.90 | 5.60 | 223.26 | 5.77 | 229.83 |
| C22:1n-9 | 0.60 | 28.56 | 0.60 | 23.80 | 0.86 | 34.14 | 0.88 | 35.04 |
| C24:1n-9 | 0.23 | 11.17 | 0.41 | 16.20 | 0.76 | 30.27 | 0.81 | 32.42 |
| **ΣMUFAs** | **34.76** | **1654.79** | **34.78** | **1383.91** | **36.65** | **1461.75** | **37.12** | **1479.32** |
| C18:2n-6 | 32.05 | 1525.96 | 31.91 | 1269.72 | 25.14 | 1002.86 | 24.46 | 974.76 |
| C18:3n-6 | 0.10 | 4.88 | 0.10 | 3.92 | 0.10 | 4.01 | <LOQ | <LOQ |
| C20:2n-6 | 0.38 | 18.24 | 0.40 | 15.86 | 0.34 | 13.44 | 0.36 | 14.28 |
| C20:3n-6 | 0.14 | 6.51 | 0.14 | 5.42 | 0.10 | 4.06 | 0.10 | 4.16 |
| C20:4n-6 (ARA) | 0.24 | 11.28 | 0.24 | 9.43 | 0.44 | 17.50 | 0.45 | 18.00 |
| C22:4n-6 | <LOQ | <LOQ | <LOQ | <LOQ | <LOQ | <LOQ | <LOQ | <LOQ |
| C22:5n-6 | 0.11 | 5.22 | 0.10 | 4.11 | 0.13 | 5.37 | 0.12 | 4.84 |
| **Σn-6 PUFAs** | **33.02** | **1572.08** | **32.88** | **1308.46** | **26.25** | **1047.25** | **25.49** | **1016.04** |
| C18:3n-3 | 4.61 | 219.56 | 4.53 | 180.11 | 2.52 | 100.34 | 2.47 | 98.48 |
| C18:4n-3 | 1.05 | 49.83 | 1.00 | 39.85 | 1.11 | 44.15 | 1.06 | 42.35 |
| C20:3n-3 | 0.14 | 6.56 | 0.17 | 6.72 | 0.14 | 5.56 | 0.17 | 6.68 |
| C20:4n-3 | 0.35 | 16.44 | 0.35 | 14.07 | 0.37 | 14.70 | 0.37 | 14.66 |
| C20:5n-3 (EPA) | 2.45 | 116.43 | 2.36 | 93.92 | 4.24 | 169.10 | 4.03 | 160.77 |
| C21:5n-3 | 0.16 | 7.46 | 0.17 | 6.67 | 0.17 | 6.77 | 0.18 | 7.11 |
| C22:5n-3 | 0.53 | 25.14 | 0.50 | 19.97 | 0.52 | 20.89 | 0.50 | 19.74 |
| C22:6n-3 (DHA) | 3.52 | 167.77 | 3.30 | 131.36 | 5.97 | 238.06 | 5.79 | 230.94 |
| **Σn-3 PUFAs** | **12.80** | **609.18** | **12.38** | **492.69** | **15.03** | **599.57** | **14.57** | **580.73** |
| C16:02 | 0.14 | 6.90 | 0.13 | 5.32 | 0.23 | 9.33 | 0.23 | 9.19 |
| C16:03 | 0.10 | 4.99 | 0.10 | 4.06 | 0.09 | 3.68 | 0.09 | 3.48 |
| C16:04 | 0.18 | 8.75 | 0.18 | 6.96 | 0.16 | 6.29 | 0.15 | 5.90 |
| | **0.43** | **20.65** | **0.41** | **16.35** | **0.48** | **19.29** | **0.47** | **18.58** |
| **ΣPUFAs** | **46.25** | **2201.92** | **45.68** | **1817.50** | **41.77** | **1666.11** | **40.53** | **1615.36** |
| **Total** | **100.00** | **4760.86** | **100.00** | **3978.96** | **100.00** | **3988.89** | **100.00** | **3985.77** |
| **n-3/n-6** | **0.39** | | **0.38** | | **0.57** | | **0.57** | |

Limit of quantification (LOQ) for fatty acid analysis is 0.06%. C18:1n-9: oleic acid; C22:1n-9: erucic acid; C18:2n-6: linoleic acid; C18:3n-3: linolenic acid; C20:4n-6 (ARA): arachidonic acid; C20:5n-3 (EPA): eicosapentaenoic acid; C22:6n-3 (DHA): docosahexaenoic acid; SFA: saturated fatty acid; MUFA: monounsaturated fatty acid; PUFA: polyunsaturated fatty acid; HUFA: highly unsaturated fatty acids.

Table S3. Amino acid analysis of *L. vannamei* fed the experimental diets for 8 weeks (g/100g). D1 (SE) = soya and pea extruded, D2 (SS) = soya and pea steamed, D3 (FE) = Fishmeal extruded and D4 (FS) = fishmeal steamed.

| | **Experimental Diets** | | | | One-way ANOVA | | Two-way ANOVA p-value | | |
|---|---|---|---|---|---|---|---|---|---|
| | **D1 (SE)** | **D2 (SS)** | **D3 (FE)** | **D4 (FS)** | *F-value* | *p-value* | **Feed processing type** | **Protein source** | Interaction |
| **Aspartic acid** | 7.03 ± 0.55 | 6.75 ± 0.37 | 7.21 ± 0.08 | 6.97 ± 0.20 | 0.878 | 0.492 | 0.237 | 0.349 | 0.923 |
| **Serine** | 2.79 ± 0.10 | 2.95 ± 0.65 | 2.64 ± 0.11 | 2.79 ± 0.14 | 0.415 | 0.747 | 0.453 | 0.453 | 0.993 |
| **Glutamic acid** | 11.00 ± 0.91 | 10.39 ± 0.63 | 11.44 ± 0.49 | 10.80 ± 0.43 | 1.379 | 0.318 | 0.130 | 0.288 | 0.976 |
| **Proline** | 3.87 ± 0.33 | 4.01 ± 0.52 | 3.71 ± 0.23 | 3.63 ± 0.32 | 0.646 | 0.607 | 0.885 | 0.235 | 0.622 |
| **Glycine** | 5.35 ± 0.03 | 5.59 ± 1.08 | 5.72 ± 0.34 | 5.85 ± 0.18 | 0.422 | 0.743 | 0.588 | 0.366 | 0.868 |
| **Alanine** | 3.78 ± 0.24 | 3.79 ± 0.50 | 3.86 ± 0.05 | 3.75 ± 0.07 | 0.074 | 0.972 | 0.773 | 0.897 | 0.743 |
| **Valine** | 3.02 ± 0.16 | 2.78 ± 0.10 | 3.09 ± 0.01 | 2.99 ± 0.17 | 3.282 | 0.079 | 0.048 | 0.097 | 0.380 |
| **Isoleucine** | 2.77 ± 0.14$^{b}$ | 2.53 ± 0.06$^{a}$ | 2.82 ± 0.01$^{b}$ | 2.72 ± 0.09$^{b}$ | 6.288 | 0.017 | 0.010 | 0.046 | 0.191 |
| **Leucine** | 4.70 ± 0.31 | 4.50 ± 0.24 | 4.86 ± 0.09 | 4.66 ± 0.17 | 1.370 | 0.320 | 0.152 | 0.241 | 0.980 |
| **Tyrosine** | 2.52 ± 0.17 | 2.47 ± 0.06 | 2.58 ± 0.10 | 2.50 ± 0.09 | 0.580 | 0.644 | 0.317 | 0.472 | 0.860 |
| **Phenylalanine** | 3.02 ± 0.19 | 2.94 ± 0.08 | 3.08 ± 0.06 | 2.98 ± 0.12 | 0.693 | 0.582 | 0.244 | 0.505 | 0.928 |
| **Histidine** | 1.70 ± 0.15 | 1.60 ± 0.05 | 1.72 ± 0.01 | 1.65 ± 0.09 | 1.106 | 0.402 | 0.134 | 0.500 | 0.852 |
| **Lysine** | 4.98 ± 0.35$^{abc}$ | 4.57 ± 0.12$^{b}$ | 5.21 ± 0.04$^{a}$ | 4.96 ± 0.16$^{c}$ | 5.123 | 0.029 | 0.022 | 0.030 | 0.506 |
| **Arginine** | 5.24 ± 0.26 | 5.07 ± 0.44 | 5.15 ± 0.12 | 5.08 ± 0.07 | 0.276 | 0.841 | 0.446 | 0.785 | 0.753 |
| **Threonine** | 3.62 ± 1.83 | 2.52 ± 0.25 | 3.77 ± 2.16 | 3.70 ± 2.24 | 0.379 | 0.774 | 0.811 | 0.430 | 0.612 |
| **Methionine** | 1.57 ± 0.08 | 1.60 ± 0.18 | 1.65 ± 0.07 | 1.59 ± 0.02 | 0.325 | 0.808 | 0.788 | 0.558 | 0.490 |
| **Cysteine** | 0.22 ± 0.05 | 0.29 ± 0.06 | 0.26 ± 0.04 | 0.29 ± 0.02 | 1.712 | 0.241 | 0.101 | 0.382 | 0.382 |

Values are means of triplicate groups and presented as mean ± SD. Values in the same row having different superscript letters are significantly different ($p < 0.05$). The lack of superscript letter indicates no significant differences among treatments.

Table S4. Fatty acid profile of *L. vannamei* fed the experimental diets for 8 weeks (% and mg/100g). D1 (SE) = soya and pea extruded, D2 (SS) = soya and pea steamed, D3 (FE) = Fishmeal extruded and D4 (FS) = fishmeal steamed.

| Fatty acid | **D1 (SE)** | | **D2 (SS)** | | **D3 (FE)** | | **D4 (FS)** | | **One-way ANOVA** | | **Two-way ANOVA p-value** | | |
|---|---|---|---|---|---|---|---|---|---|---|---|---|---|
| | **%** | **mg. 100g$^{-1}$** | **%** | **mg. 100g$^{-1}$** | **%** | **mg. 100g$^{-1}$** | **%** | **mg. 100g$^{-1}$** | **F-value** | **p-value** | **Feed processing type** | **Protein source** | **Interaction** |
| C14:0 | 0.50 ± 0.09 | 13.92 ± 1.58 | 0.55 ±0.08 | 15.08 ±2.67 | 0.58 ±0.08 | 15.74 ±2.32 | 0.56 ± 0.10 | 16.39 ±5.17 | 0.508 | 0.687 | 0.750 | 0.351 | 0.528 |
| C15:0 | 0.33 ± 0.05 | 9.66 ± 2.77 | 0.38 ±0.02 | 10.45 ±1.48 | 0.39 ±0.03 | 10.78 ±0.75 | 0.37 ±0.02 | 10.62 ±1.26 | 1.747 | 0.235 | 0.616 | 0.201 | 0.120 |
| C16:0 | 16.80 ± 0.64[a] | 477.36 ± 80.07 | 16.93 ±0.11[a] | 468.62 ±39.16 | 18.51 ±0.45[b] | 503.77 ±7.42 | 17.77±0.29[ab] | 511.82 ±79.44 | 10.587 | 0.004 | 0.250 | $8.23 \times 10^{-4}$ | 0.112 |
| C18:0 | 8.33 ± 0.40 | 234.94 ± 19.35 | 8.48 ±0.40 | 233.95 ±7.47 | 8.55 ±0.14 | 232.79 ±2.85 | 8.54 ±0.62 | 245.20 ±35.01 | 0.179 | 0.908 | 0.792 | 0.573 | 0.742 |
| C20:0 | 0.47 ± 0.03 | 13.35 ± 1.14 | 0.50 ±0.03 | 13.73 ±0.87 | 0.44 ±0.03 | 11.85 ±0.99 | 0.47 ±0.03 | 13.40 ±1.18 | 2.206 | 0.165 | 0.124 | 0.093 | 0.854 |
| C22:0 | 0.61 ± 0.09 | 17.20 ± 0.86 | 0.66 ±0.08 | 18.16 ±1.46 | 0.58 ±0.05 | 15.95 ±1.44 | 0.76 ±0.12 | 21.86 ±3.42 | 2.479 | 0.135 | 0.051 | 0.505 | 0.231 |
| C24:0 | 0.24 ± 0.04 | 6.75 ± 0.59 | 0.26 ±0.03 | 7.16 ±0.43 | 0.26 ±0.01 | 7.02 ±0.21 | 0.34 ±0.06 | 9.83 ±1.89 | 3.761 | 0.060 | 0.056 | 0.070 | 0.201 |
| **ΣSFAs** | **27.29 ± 0.45 [a]** | **773.18 ± 103.60** | **27.75 ±0.40 [ac]** | **767.16 ±50.16** | **29.31 ±0.37 [b]** | **797.89 ±0.40** | **28.81±0.85[bc]** | **829.12 ±120.56** | **8.533** | **0.007** | **0.956** | **$1.29 \times 10^{-3}$** | **0.175** |
| C16:1n-9 | 0.25 ± 0.05 | 7.10 ±2.23 | 0.23 ±0.02 | 6.28 ±0.77 | 0.23 ±0.01 | 6.13 ±0.20 | 0.24 ±0.04 | 6.87 ±1.79 | 0.237 | 0.868 | 0.806 | 0.806 | 0.467 |
| C16:1n-7 | 1.01 ± 0.06 [a] | 28.44 ±2.43 | 1.00 ±0.09 [a] | 27.85 ±4.45 | 1.44 ±0.07 [b] | 39.26 ±2.33 | 1.40 ±0.19 [b] | 40.65 ±10.72 | 12.387 | $2.24 \times 10^{-3}$ | 0.740 | $2.95 \times 10^{-4}$ | 0.812 |
| C18:1n-9 | 17.36 ±0.86 | 493.78 ±89.32 | 17.22 ±0.78 | 477.43 ±56.85 | 16.06 ±0.69 | 437.53 ±24.22 | 15.30 ±1.06 | 440.89 ±78.15 | 3.918 | 0.054 | 0.390 | 0.012 | 0.545 |
| C18:1n-7 | 2.28 ±0.09[a] | 64.73 ±9.16 | 2.19 ±0.06 [a] | 60.56 ±6.05 | 3.11 ±0.01[b] | 84.69 ±1.01 | 2.92 ±0.16[b] | 84.19 ±15.37 | 65.881 | $5.54 \times 10^{-6}$ | 0.033 | $7.36 \times 10^{-7}$ | 0.417 |
| C20:1n-11 | 0.24 ±0.04[a] | 7.05 ±2.23 | 0.23 ±0.04[a] | 6.44 ±1.38 | 0.44 ±0.03[b] | 12.03 ±0.48 | 0.35 ±0.08[ab] | 10.13 ±3.42 | 12.432 | 0.002 | 0.097 | $4.88 \times 10^{-4}$ | 0.196 |
| C20:1n-9 | 2.17 ±0.17[a] | 61.72 ±12.55 | 2.07 ±0.09[a] | 57.26 ±6.59 | 2.65 ±0.05[b] | 72.14 ±1.25 | 2.33 ±0.33[ab] | 67.62 ±17.68 | 5.393 | 0.025 | 0.095 | 0.009 | 0.354 |
| C20:1n-7 | 0.11 ±0.01 | 3.17 ±0.76 | 0.11 ±0.01 | 3.21 ±0.63 | 0.16 ±0.02 | 4.37 ±0.63 | 0.12 ±0.04 | 3.43 ±1.38 | 3.794 | 0.058 | 0.132 | 0.059 | 0.089 |

| | | | | | | | | | | | | | |
|---|---|---|---|---|---|---|---|---|---|---|---|---|---|
| C22:1n-11 | 0.94 ±0.21 | 26.99 ±9.29 | 0.89 ±0.09 | 24.66 ±4.19 | 1.32 ±0.05 | 35.98 ±1.02 | 0.85 ±0.43 | 25.22 ±15.80 | 2.365 | 0.147 | 0.104 | 0.248 | 0.177 |
| C22:1n-9 | 0.21 ±0.04 | 5.99 ±1.88 | 0.21 ±0.02 | 5.75 ±1.05 | 0.26 ±0.01 | 6.99 ±0.13 | 0.22 ±0.04 | 6.28 ±1.86 | 1.872 | 0.213 | 0.284 | 0.123 | 0.284 |
| C24:1n-9 | 0.43 ±0.09 | 11.93 ±1.20 | 0.45 ±0.04 | 12.45 ±0.03 | 0.50 ±0.02 | 13.69 ±0.37 | 0.58 ±0.07 | 16.53 ±2.58 | 3.718 | 0.061 | 0.207 | 0.019 | 0.455 |
| **ΣMUFAs** | **25.00 ±1.23** | **710.89±127.80** | **24.59 ±1.10** | **681.88 ±81.07** | **26.17 ±0.65** | **712.81 ±26.52** | **24.30 ±2.18** | **701.81 ±144.66** | **1.027** | **0.431** | **0.199** | **0.601** | **0.392** |
| C18:2n-6 | 17.50 ±1.50$^{a}$ | 499.26 ±107.40 | 17.33 ±0.65$^{a}$ | 480.14 ±51.32 | 13.45 ±0.91$^{b}$ | 366.08 ±21.79 | 13.29 ±0.30$^{b}$ | 382.67 ±59.36 | 18.280 | $6.12\times10^{-4}$ | 0.773 | $7.62\times10^{-5}$ | 0.995 |
| C18:3n-6 | 0.22 ±0.02$^{ab}$ | 6.37 ±1.19 | 0.24 ±0.02$^{b}$ | 6.57 ±0.74 | 0.20 ±0.01$^{ad}$ | 5.38 ±0.13 | 0.18 ±0.01$^{cd}$ | 5.28 ±0.82 | 11.222 | 0.003 | 0.833 | $6.04\times10^{-4}$ | 0.085 |
| C20:2n-6 | 2.23 ±0.26$^{a}$ | 63.61 ±14.67 | 2.18 ±0.8$^{a}$ | 60.28 ±4.41 | 1.52 ±0.08$^{b}$ | 41.38 ±1.64 | 1.49 ±0.04$^{b}$ | 42.82 ±6.18 | 24.016 | $2.35\times10^{-4}$ | 0.643 | $2.87\times10^{-5}$ | 0.938 |
| C20:3n-6 | 0.09 ±0.01$^{a}$ | 2.58 ±0.52 | 0.08 ±0.01$^{a}$ | 2.41 ±0.34 | 0.00$^{b}$ | 0 | 0.02 ±0.00$^{b}$ | 2.06 ±0.00 | 18.300 | $6.10\times10^{-4}$ | 0.545 | $8.60\times10^{-5}$ | 0.242 |
| C20:4n-6 (ARA) | 2.06 ±0.39 | 57.39 ±4.87 | 2.01 ±0.14 | 55.29 ±0.96 | 2.14 ±0.16 | 58.32 ±5.16 | 2.45 ±0.38 | 69.88 ±10.90 | 1.359 | 0.323 | 0.480 | 0.159 | 0.321 |
| C22:4n-6 | 0.07 ±0.00 | 1.79 ±0.00 | <LOQ | <LOQ | <LOQ | <LOQ | <LOQ | <LOQ | | | | | |
| C22:5n-6 | 0.25 ±0.05 | 7.08 ±1.01 | 0.25 ±0.03 | 6.91 ±0.43 | 0.21 ±0.01 | 5.70 ±0.07 | 0.24 ±0.02 | 6.91 ±0.50 | 0.952 | 0.460 | 0.496 | 0.248 | 0.399 |
| **Σn-6 PUFAs** | **22.38 ±1.40$^{a}$** | **636.90 ±119.57** | **22.09 ±0.55$^{a}$** | **611.60 ±56.56** | **17.52 ±0.84$^{b}$** | **476.87 ±18.80** | **17.67 ±0.10$^{b}$** | **508.24 ±71.49** | **29.099** | **$1.18\times10^{-4}$** | **0.892** | **$1.41\times10^{-5}$** | **0.675** |
| C18:3n-3 | 1.34 ±0.01$^{a}$ | 38.08 ±4.87 | 1.38 ±0.08$^{a}$ | 38.14 ±3.97 | 0.83 ±0.11$^{b}$ | 22.69 ±3.22 | 0.78 ±0.15 $^{b}$ | 22.52 ±5.70 | 31.194 | $9.16\times10^{-5}$ | 0.888 | $1.11\times10^{-5}$ | 0.457 |
| C18:4n-3 | 0.16 ±0.03 | 4.43 ±0.30 | 0.16 ±0.02 | 4.37 ±0.56 | 0.16 ±0.04 | 4.40 ±1.05 | 0.15 ±0.06 | 4.44 ±1.88 | 0.021 | 0.995 | 0.825 | 0.941 | 0.941 |
| C20:3n-3 | 0.26 ±0.04$^{a}$ | 7.52 ±1.94 | 0.25 ±0.01$^{a}$ | 6.78 ±0.71 | 0.16 ±0.01$^{b}$ | 4.35 ±0.11 | 0.15 ±0.01$^{b}$ | 4.17 ±0.58 | 27.933 | $1.36\times10^{-4}$ | 0.327 | $1.71\times10^{-5}$ | 0.885 |
| C20:4n-3 | 0.22 ±0.02$^{a}$ | 6.21 ±1.11 | 0.21 ±0.01$^{ab}$ | 5.82 ±0.82 | 0.18 ±0.01$^{ab}$ | 5.01 ±0.07 | 0.18 ±0.03$^{b}$ | 5.02 ±1.24 | 4.865 | 0.033 | 0.403 | 0.006 | 0.864 |
| C20:5n-3 (EPA) | 9.77 ±1.33$^{a}$ | 273.51 ±4.26 | 9.85 ±0.45$^{a}$ | 272.04 ±11.06 | 11.06 ±0.22$^{ab}$ | 301.25 ±9.30 | 12.11 ±0.88$^{b}$ | 346.56 ±32.64 | 5.302 | 0.026 | 0.274 | 0.006 | 0.346 |
| C21:5n-3 | 0.13 ±0.01 | 3.57 ±0.46 | 0.12 ±0.01 | 3.51 ±0.42 | 0.11 ±0.01 | 2.93 ±0.13 | 0.11 ±0.02 | 3.07 ±0.73 | 3.154 | 0.086 | 0.789 | 0.016 | 0.789 |
| C22:5n-3 | 0.60 ±0.04$^{a}$ | 16.99 ±2.13 | 0.59 ±0.02$^{a}$ | 16.45 ±1.88 | 0.45 ±0.02$^{b}$ | 12.09 ±0.53 | 0.45 ±0.03$^{b}$ | 13.14 ±2.34 | 26.128 | $1.74\times10^{-4}$ | 1.000 | $2.10\times10^{-5}$ | 0.698 |

| | | | | | | | | | | | | | |
|---|---|---|---|---|---|---|---|---|---|---|---|---|---|
| C22:6n-3 (DHA) | 8.07 ±0.81[a] | 226.73 ±9.51 | 7.95 ±0.44[a] | 219.39 ±11.82 | 9.60 ±0.04[b] | 261.45 ±3.45 | 10.14 ±0.34[b] | 291.39 ±37.94 | 15.044 | $1.18\times10^{-3}$ | 0.483 | $1.74\times10^{-4}$ | 0.271 |
| **Σn-3 PUFAs** | **20.55 ±2.10**[a] | **577.03 ±14.53** | **20.51 ±0.88**[a] | **566.50 ±30.42** | **22.56 ±0.39**[ab] | **614.17 ±17.54** | **24.07 ±0.98**[b] | **690.30 ±75.89** | **5.638** | **0.023** | **0.340** | **0.005** | **0.314** |
| **ΣPUFAs** | **42.93 ±1.05**[a] | **1213.93 ±133.54** | **42.60 ±0.39**[a] | **1178.10 ±81.34** | **40.07 ±0.46**[b] | **1091.04 ±10.95** | **41.74 ±1.04**[ab] | **1198.54 ±146.55** | **7.681** | **0.010** | **0.185** | **0.004** | **0.062** |
| 16:0 DMA | 1.66 ±0.15 | 46.64 ±2.23 | 1.75 ±0.10 | 48.27 ±1.43 | 1.66 ±0.05 | 45.20 ±1.85 | 1.89 ±0.20 | 54.01 ±4.82 | 1.938 | 0.202 | 0.075 | 0.385 | 0.406 |
| 18:0 DMA | 3.12 ±0.29 | 87.74 ±4.67 | 3.31 ±0.22 | 91.24 ±1.57 | 2.78 ±0.13 | 75.80 ±3.86 | 3.25 ±0.35 | 92.96 ±10.84 | 2.472 | 0.136 | 0.060 | 0.222 | 0.383 |
| **ΣDMAs** | **4.78 ±0.44** | **134.38 ±6.89** | **5.06 ±0.32** | **139.51 ±2.99** | **4.44 ±0.18** | **121.01 ±5.57** | **5.15 ±0.55** | **146.97 ±15.66** | **1.907** | **0.207** | **0.065** | **0.605** | **0.379** |
| **Total** | **100.00 ±0.00** | **2832.39 ±367.33** | **100.00 ±0.00** | **2766.65 ±214.46** | **100.00 ±0.00** | **2722.74 ±34.92** | **100** | **2876.43 ±404.71** | | | | | |

Values are means of triplicate groups and presented as mean ± SD. Values of the percentages in the same row having different superscript letters are significantly different ($p < 0.05$). The lack of superscript letter near the percent values indicates no significant differences among treatments. SFAs (Saturated Fatty Acids), MUFAs (Monounsaturated Fatty Acids), n-6 PUFAs (Omega-6 Polyunsaturated Fatty Acids), n-3 PUFAs (Omega-3 Polyunsaturated Fatty Acids), DMAs (Dimethyl Acids).
